# A machine learning framework for interpreting phylogenetic tree patterns in interkingdom horizontal gene transfer

**DOI:** 10.64898/2026.05.21.726852

**Authors:** Kevin Aguirre-Carvajal, Vinicio Armijos-Jaramillo, Cristian R. Munteanu

**Affiliations:** Department of Computer Science and Information Technologies, Faculty of Computer Science, CITIC Research Center of Information and Communication Technologies, University of A Coruña, Campus Elviña s/n, 15071 A Coruña, Spain; Bio-Cheminformatics Research Group, Universidad de Las Américas, Quito 170513, Ecuador; Carrera de Ingeniería en Biotecnología, Facultad de Ingeniería y Ciencias Aplicadas, Universidad de Las Américas, Quito 170513, Ecuador

**Keywords:** interkingdom horizontal gene transfer, phylogenomics, machine learning, phylogenetic pattern classification, computational evolutionary biology

## Abstract

Horizontal gene transfer (HGT), the movement of genetic material between unrelated organisms, is widely recognized as an important driver of genome evolution in bacteria. In eukaryotes, however, the evolutionary impact of HGT remains debated. The identification of interkingdom HGT (iHGT) is especially challenging due to the lack of gold standard methods.

Traditionally, iHGT identification has relied on manual inspection of phylogenetic trees, a process that is subjective, difficult to reproduce, and not scalable to large datasets. In this study, we present a computational framework that formalizes phylogenetic tree interpretation into a supervised machine-learning problem. We define five recurrent phylogenetic patterns—iHGT, NoHGT, Limited donor evidence, Multiple major clades (Multiple MC), and Patchy phylogeny—capturing clear and ambiguous evolutionary scenarios.

To operationalize these patterns, we developed a feature-extraction pipeline that quantifies taxonomic composition and phylogenetic topology using seven biological descriptors derived from gene trees. These features were used to train and evaluate multiple machine-learning models, among which a Random Forest (RF) classifier achieved the best performance (AUC–ROC = 0.98; accuracy = 0.89). Model interpretability analyses revealed that topological distance to additional clades and lineage diversity are the most informative predictors, reflecting key signals used in expert-driven phylogenetic interpretation.

The RF model was further validated using 1,000 simulated phylogenies and 1,438 real iHGT candidates, achieving low misclassification rates (7.8% and 10.43%, respectively). Benchmarking against AVP (Alienness vs. Predictor), a comparable tool for iHGT detection, demonstrated improved performance across all evaluation metrics, highlighting the advantages of incorporating global phylogenetic structure into the classification process. This study provides a reproducible and scalable framework for phylogenetic pattern classification that captures complex evolutionary signals while maintaining biological interpretability. Beyond improving iHGT detection, the approach offers a more nuanced representation of evolutionary scenarios by explicitly accounting for inconclusive cases, supporting more robust inference in comparative genomics.

**Authors summary:** Horizontal gene transfer is the movement of genes between unrelated organisms rather than through normal inheritance from parent to offspring. While this process is known to play a major role in bacterial evolution, its importance in complex organisms such as fungi, plants, and animals remains debated. One reason for this uncertainty is that identifying these events often depends on manually interpreting phylogenetic trees, a process that can be subjective, difficult to reproduce, and impractical for analyzing the rapidly growing amount of genomic data.

In this study, we developed a computational framework that transforms phylogenetic tree interpretation into a machine-learning problem. Instead of simply classifying genes as transferred or non-transferred, our approach recognizes several distinct evolutionary scenarios, including cases where the evidence is ambiguous or inconclusive. To achieve this, we extracted biologically meaningful features from phylogenetic trees describing evolutionary relationships and taxonomic diversity, and used them to train machine-learning models capable of recognizing recurrent phylogenetic patterns.

Our framework successfully classified complex evolutionary scenarios and outperformed an existing automated method for interkingdom horizontal gene transfer detection. More broadly, this work demonstrates how expert-driven evolutionary reasoning can be translated into scalable and reproducible computational approaches. As genomic datasets continue to expand, such methods may help improve evolutionary inference and support more rigorous comparative genomics analyses.

## Introduction

Horizontal gene transfer (HGT) refers to the movement of genetic material between non-related organisms, bypassing vertical inheritance. In bacteria, HGT is widely recognized as a major driver of genetic innovation and diversification. In contrast, its prevalence and evolutionary impact in eukaryotes remain the subject of ongoing debate. Particularly contentious is the occurrence of interdomain horizontal gene transfer (iHGT), which involves gene transfer between organisms from different biological domains. Given the profound genetic divergence separating these lineages, such events would be expected to be rare or even negligible. Nevertheless, an increasing number of studies have reported putative cases of iHGT in eukaryotes (1–4).

At the same time, recent analyses have suggested that many reported iHGT cases may be better explained by alternative evolutionary scenarios, and that methodological biases in detection approaches can substantially influence both the number and the nature of inferred candidates (5–8).

In this context, standardized, replicable, and biologically grounded approaches are required to improve confidence in iHGT detection and to more accurately assess its evolutionary impact in eukaryotes. Two principal methodological frameworks are commonly employed to identify HGT candidates: parametric and phylogenetic approaches.

Parametric methods rely on genome-specific compositional signatures, such as GC content, third-codon position bias, codon usage, amino acid frequencies, and k-mer distributions, among others (9). These approaches have been applied predominantly in bacterial systems (10–13) and are based on detecting discrepancies between the compositional features of the putative donor fragment and those of the recipient genome. However, this strategy presents several limitations. Intrinsic heterogeneity within recipient genomes can lead to overestimation of HGT-derived segments. Moreover, transferred sequences undergo amelioration over time, progressively converging toward the compositional profile of the host genome. Consequently, parametric approaches have limited power to detect ancient or distantly related HGT events, like interdomain lateral transferences.

Several tools implement this strategy, including SigHunter (14), TF-IDF (15), IslandPath-DIMOB (16), HGT-DB (17), Islander (11), and NECK (13). More recently, synteny-based analyses have been incorporated to infer HGT events between closely related bacteria (18,19), with the aim of mitigating some of the compositional biases described above.

In contrast, phylogenetic approaches exploit the evolutionary signal contained in gene or protein sequences to identify incongruences with the expected species phylogeny. Such conflicts are typically detected through explicit phylogenetic reconstructions or through surrogate, implicit methods that estimate evolutionary distances or sequence similarity among genes (20).

Implicit methods infer phylogenetic relationships without reconstructing explicit trees, enabling rapid screening for HGT candidates in large-scale datasets. These approaches typically rely on sequence similarity metrics and heuristic searches, which makes them computationally efficient. However, they can produce misleading inferences, as the most similar sequences identified through tools such as BLAST are not necessarily the closest evolutionary relatives in a robust phylogenetic framework (21).

Several tools have been developed based on this strategy, including DarkHorse (22), HGTector (23), HGTFinder (24), Alienness (25), BLAST2HGT (26), and ShadowCaster (27).

Explicit phylogenetic methods can be broadly divided into two categories. The first involves direct inspection and analysis of reconstructed gene trees to infer HGT events or alternative evolutionary scenarios. This approach is commonly applied when researchers reconstruct phylogenies for candidates initially identified through implicit methods, or when investigating specific genes exhibiting unusual phylogenetic patterns. However, relatively few tools are available for the automated detection of HGT candidates through direct tree analysis; AvP (28) is among the limited number of programs designed for this purpose.

The second category comprises tree reconciliation approaches, which compare gene phylogenies with a reference species tree. In principle, this framework has the capacity to discriminate among distinct evolutionary processes, including HGT, gene duplication and loss, and incomplete lineage sorting. Programs such as AnGST (29), HGTree (30), and RANGER-DTL (31) implement this strategy. Nevertheless, analyses in fungi by Dupont and Cox (2017) indicate that reconciliation methods may exhibit limited sensitivity in distinguishing artifactual HGT candidates from vertically inherited genes. Although biologically coherent in theory, the broader application of these tools requires further validation under diverse evolutionary scenarios and, at present, may not be well suited for large-scale analyses such as whole-genome datasets.

Beyond the currently available methods for detecting HGT, most tools rely on implicit assumptions about how HGT events are expected to manifest in genomic data. However, many of these expected patterns may also arise from alternative evolutionary processes or from incomplete taxonomic sampling in public databases. For instance, implicit similarity-based approaches assume that a transferred gene should exhibit greater similarity to sequences from the putative donor lineage than to those from closely related species of the recipient lineage. Yet, this pattern may also result from the absence of unsampled or unsequenced homologs closely related to the recipient lineage.

Explicit phylogenetic approaches, in turn, often depend on the experience and interpretative criteria of the researcher when identifying incongruent topologies. This subjectivity can lead to either overestimation or underestimation of robust candidates, particularly in the context of iHGT events.

Given the methodological challenges associated with HGT detection—and the limited number of approaches specifically designed to identify iHGT—we propose a direct and systematic evaluation of phylogenetic trees (an explicit approach independent of tree reconciliation) to classify candidates under alternative evolutionary scenarios. To achieve this, we aim to train machine learning models using curated phylogenetic trees that have been previously filtered and carefully annotated to distinguish iHGT candidates from other plausible evolutionary explanations. With this approach, we aim to emulate expert-driven criteria for distinguishing among iHGT candidates using explicit phylogenetic information, thereby standardizing and objectifying the detection of HGT events. We anticipate that the development of tools grounded in strict and biologically meaningful criteria will contribute to a more accurate assessment of the true evolutionary impact of HGT across diverse groups of organisms.

## Results

### Model analysis

A feature-extraction pipeline was developed to derive quantitative descriptors from phylogenetic trees. The features extracted from the training dataset is provided in Supplementary Table 6. These features were subsequently used to train multiple machine-learning models to classify phylogenetic patterns.

The predictive performance of the evaluated models is summarized in Table 1. Overall, the Random forest (RF) model achieved the best performance across most evaluation metrics, obtaining the highest AUC-ROC (0.9806 ± 0.0051), precision (0.8894 ± 0.019), recall (0.8891 ± 0.0186), F1-score (0.8878 ± 0.019), and accuracy (0.8891 ± 0.0186). According to pairwise comparisons Nemenyi test, RF showed significantly better performance than several alternative models for most metrics.

**Table 1.**
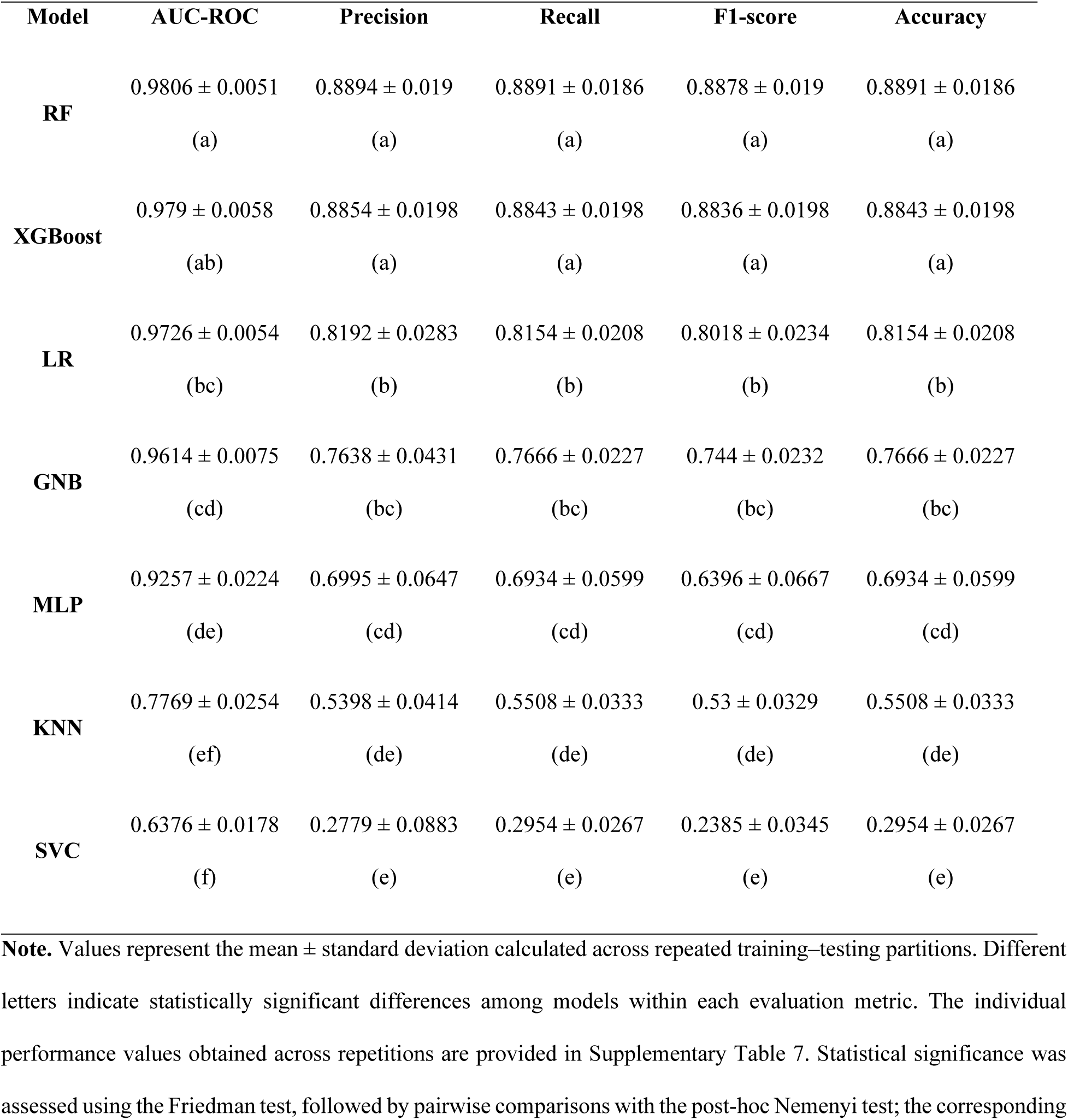
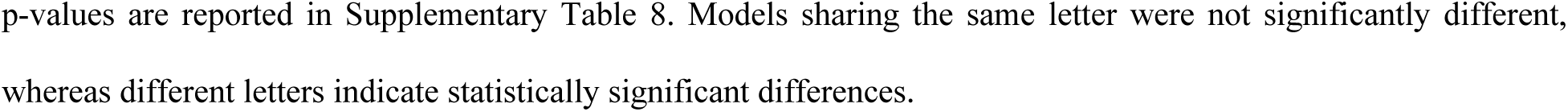
Performance comparison of machine-learning models trained to classify phylogenetic patterns.

| Model | AUC-ROC | Precision | Recall | F1-score | Accuracy |
| --- | --- | --- | --- | --- | --- |
| RF | 0.9806 ± 0.0051 | 0.8894 ± 0.019 | 0.8891 ± 0.0186 | 0.8878 ± 0.019 | 0.8891 ± 0.0186 |
|  | (a) | (a) | (a) | (a) | (a) |
| XGBoost | 0.979 ± 0.0058 | 0.8854 ± 0.0198 | 0.8843 ± 0.0198 | 0.8836 ± 0.0198 | 0.8843 ± 0.0198 |
|  | (ab) | (a) | (a) | (a) | (a) |
| LR | 0.9726 ± 0.0054 | 0.8192 ± 0.0283 | 0.8154 ± 0.0208 | 0.8018 ± 0.0234 | 0.8154 ± 0.0208 |
|  | (bc) | (b) | (b) | (b) | (b) |
| GNB | 0.9614 ± 0.0075 | 0.7638 ± 0.0431 | 0.7666 ± 0.0227 | 0.744 ± 0.0232 | 0.7666 ± 0.0227 |
|  | (cd) | (bc) | (bc) | (bc) | (bc) |
| MLP | 0.9257 ± 0.0224 | 0.6995 ± 0.0647 | 0.6934 ± 0.0599 | 0.6396 ± 0.0667 | 0.6934 ± 0.0599 |
|  | (de) | (cd) | (cd) | (cd) | (cd) |
| KNN | 0.7769 ± 0.0254 | 0.5398 ± 0.0414 | 0.5508 ± 0.0333 | 0.53 ± 0.0329 | 0.5508 ± 0.0333 |
|  | (ef) | (de) | (de) | (de) | (de) |
| SVC | 0.6376 ± 0.0178 | 0.2779 ± 0.0883 | 0.2954 ± 0.0267 | 0.2385 ± 0.0345 | 0.2954 ± 0.0267 |
|  | (f) | (e) | (e) | (e) | (e) |

**Note.** Values represent the mean ± standard deviation calculated across repeated training–testing partitions. Different letters indicate statistically significant differences among models within each evaluation metric. The individual performance values obtained across repetitions are provided in Supplementary Table 7. Statistical significance was assessed using the Friedman test, followed by pairwise comparisons with the post-hoc Nemenyi test; the corresponding
p-values are reported in Supplementary Table 8. Models sharing the same letter were not significantly different, whereas different letters indicate statistically significant differences.

The XGBoost model showed performance comparable to RF for precision, recall, F1-score, and accuracy, although it exhibited slightly lower values for AUC-ROC. LR and GNB models achieved intermediate performance, with relatively high AUC-ROC values but lower precision and recall compared with RF and XGBoost. In contrast, MLP model showed moderate performance across most metrics, while the KNN and SVC models exhibited the lowest predictive performance.

Although RF and XGBoost showed statistically comparable performance for some metrics, RF consistently achieved the highest mean values across all metrics. RF was therefore selected due to its robustness and greater interpretability, particularly through stable feature importance estimates that facilitate the identification of phylogenetic features driving classification.

The RF model was further optimized using two hyperparameter tuning strategies: randomized search and grid search. The optimal parameter configurations obtained for each strategy, as well as those of the baseline model, are provided in Supplementary Table 9.

Pairwise comparisons using the Nemenyi post-hoc test indicated that the RF model optimized through grid search performed similarly to the baseline RF model across most evaluation metrics, with the exception of AUC–ROC, for which the baseline RF model performed significantly better (AUC–ROC: *p* = 0.014; Precision: *p* = 0.9155; Recall: *p* = 0.9986; F1-score: *p* = 0.9516; Accuracy: *p* = 0.9986).

In contrast, the RF model optimized using randomized search performed significantly worse than the baseline RF model for several metrics, including precision, recall, F1-score, and accuracy, while no significant difference was observed for AUC–ROC (AUC–ROC: *p* = 0.9781; Precision: *p* = 0.0039; Recall: *p* = 0.0011; F1-score: *p* = 0.0027; Accuracy: *p* = 0.0011).

Despite exploring a wide range of parameter combinations, no significant improvement in predictive performance was observed across evaluation metrics. Based on these results, the baseline RF model was retained for subsequent implementation. The repeated performance values used for the statistical analyses are provided in Supplementary Table 7.

Once the best-performing model was selected, its class-specific discriminatory performance was further evaluated using a multiclass receiver operating characteristic (ROC) analysis (Figure 1). The RF classifier showed excellent discriminative ability across all phylogenetic categories.

**Figure 1.**
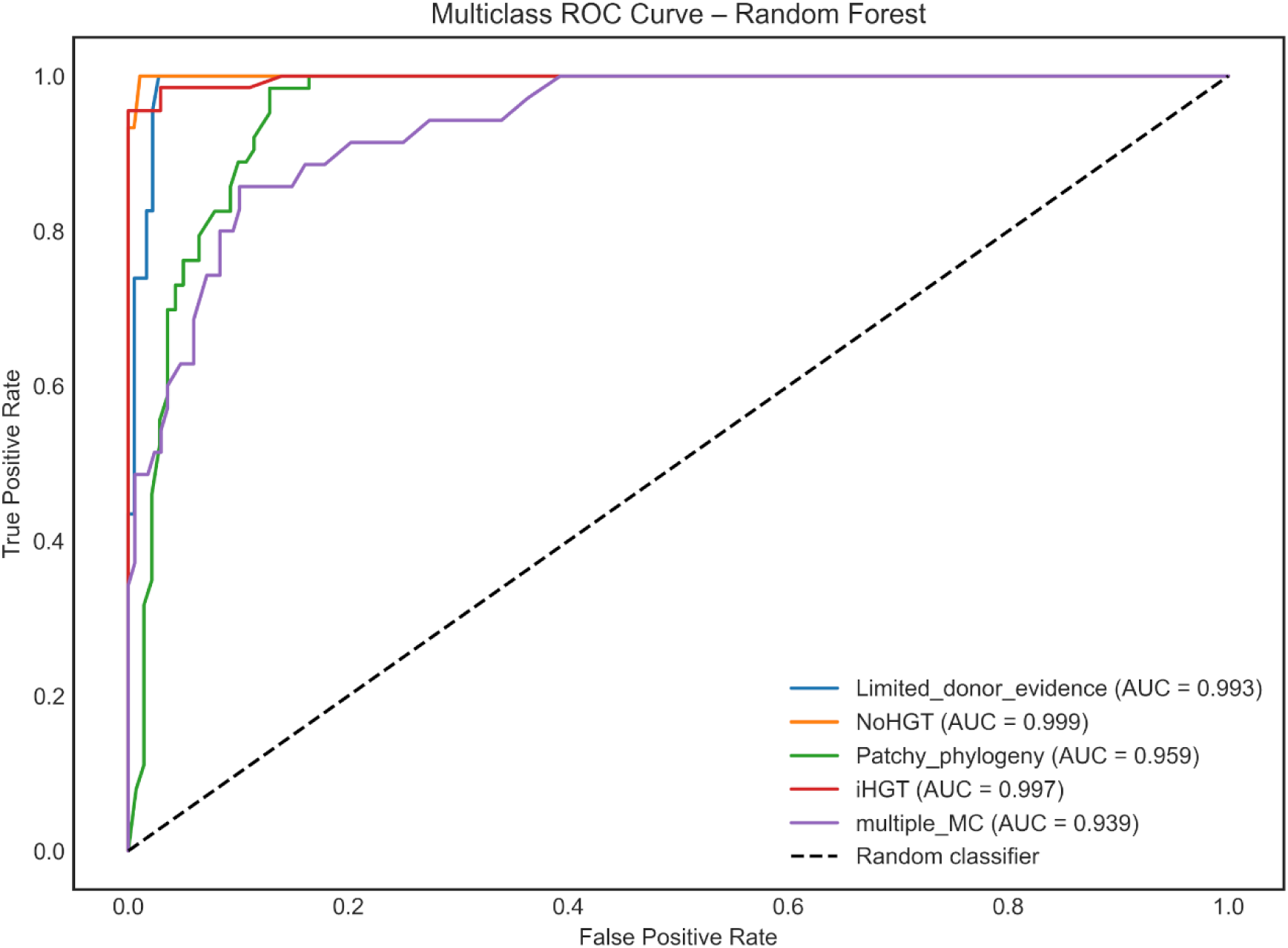
Multiclass receiver operating characteristic (ROC) curves illustrating the discriminatory performance of the Random Forest model for each phylogenetic category using a one-vs-rest approach. The area under the curve (AUC) values indicate excellent classification performance across all classes, ranging from 0.939 to 0.999. The dashed diagonal line represents the expected performance of a random classifier (AUC = 0.5).

The area under the curve (AUC) values ranged from 0.939 to 0.999, indicating excellent classification performance. The model achieved an AUC of 0.999 for the NoHGT and 0.997 for iHGT classes, demonstrating excellent separation between these categories and the remaining classes in the test partitions. Similarly high performance was observed for Limited donor evidence (AUC = 0.993), reflecting strong predictive accuracy for this phylogenetic pattern.

Slightly lower, yet still very high, performance was observed for the Patchy phylogeny (AUC = 0.959) and Multiple MC (AUC = 0.939) classes, suggesting that these phylogenetic patterns are comparatively more challenging for the model to distinguish.

To further investigate which variables contributed most to the predictive performance of the model, a permutation feature importance analysis was conducted using the decrease in weighted multiclass AUC as the evaluation metric (Figure 2). This approach quantifies the reduction in model performance after randomly permuting the values of each feature in the test set, thereby estimating its contribution to classification accuracy.

**Figure 2.**
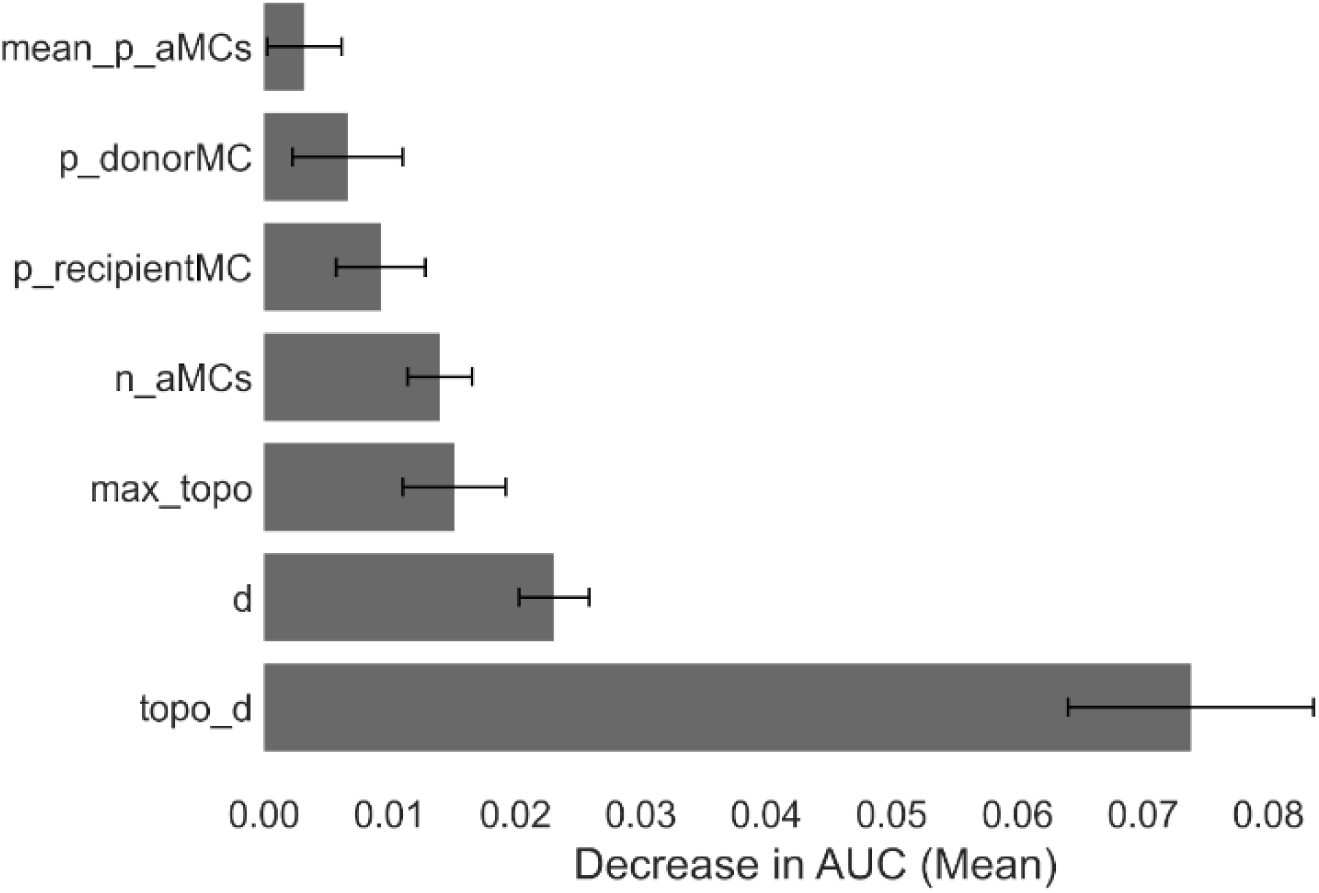
Permutation feature importance analysis for the Random Forest model. Feature importance was estimated by measuring the mean decrease in weighted multiclass AUC after randomly permuting each feature in the test dataset. Bars represent the mean decrease in AUC, and error bars indicate the standard deviation across permutations. Larger decreases indicate greater importance of the corresponding feature for model predictions.

The results indicate that *topo_d* was by far the most influential feature, producing the largest decrease in AUC when permuted (mean decrease ≈ 0.07). This value was substantially higher than that observed for any other feature, highlighting the dominant role of this topological metric in the classification of phylogenetic patterns.

A second group of features exhibited moderate importance, including *d, max_topo, n_uMCs p_donorMC* and *p_receptorMC*, all of which produced smaller but consistent decreases in AUC. These variables capture aspects of the distribution of sequences across major clades and topological relationships within the tree, suggesting that the model relies on both structural and proportional signals derived from phylogenetic topology.

In contrast, features such as *mean_p_uMCs* showed relatively minor contributions, as permuting these variables resulted in only marginal decreases in AUC. The low importance of this variable indicates that they provide limited additional predictive information beyond the more influential topological and clade-distribution features.

Additionally, to interpret the contribution of individual features to the classification process, a SHAP (SHapley Additive exPlanations) analysis was performed on the best model. The results (Figure 3) show the mean absolute SHAP values for the most influential features across all predicted classes, representing their average impact on the model output.

**Figure 3.**
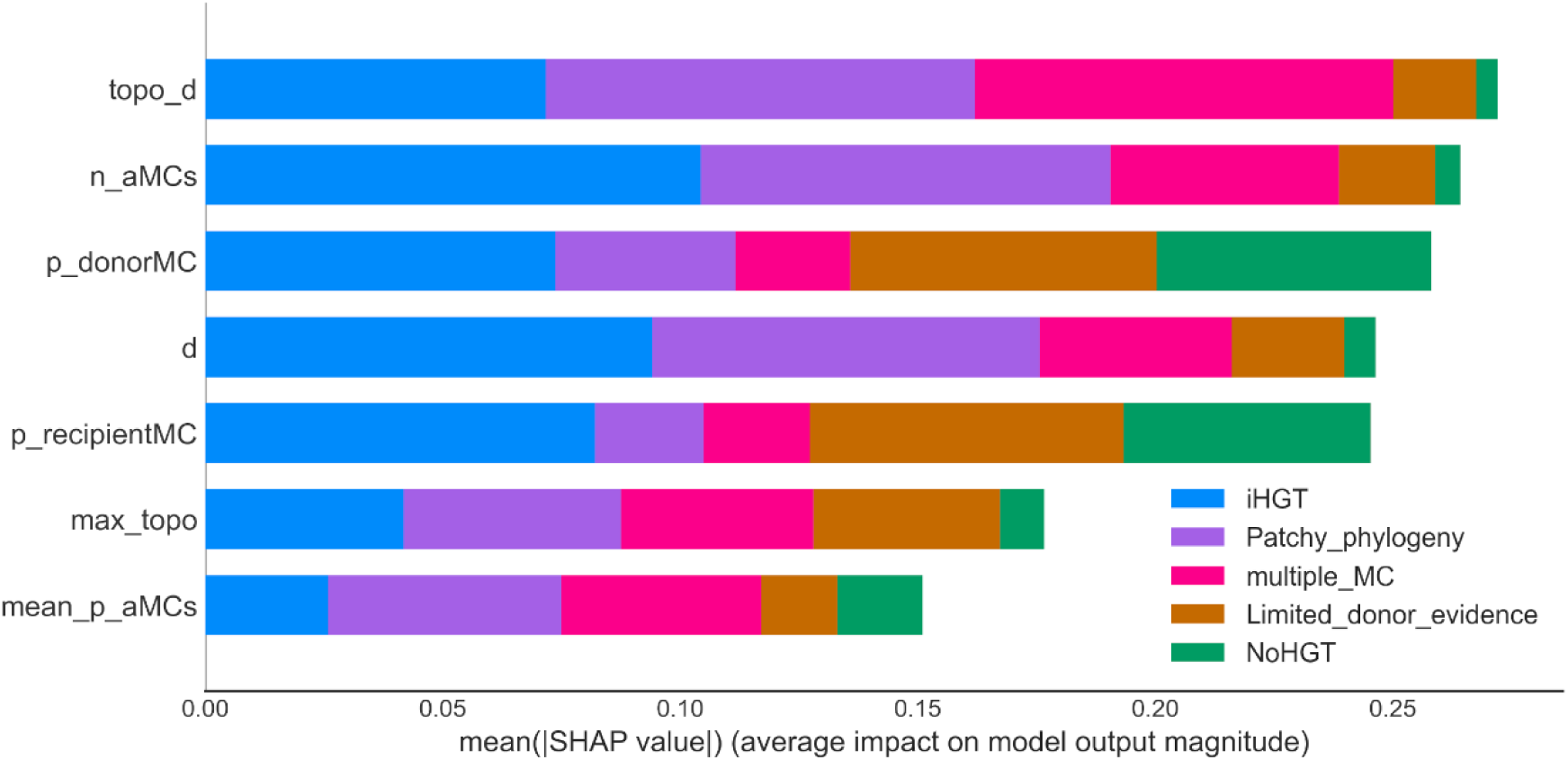
SHAP-based feature importance for the Random Forest classifier. Bars represent the mean absolute SHAP value for each feature across all samples, indicating their average contribution to the model predictions. Colors within each bar correspond to the relative contribution of each phylogenetic class. Higher SHAP values indicate a greater influence of the feature on the model output.

Among the evaluated features, the topological distance between the query sequence and the closest sequence belonging to an additional major clade (*topo_d*) emerged as the most influential predictor, followed by the number of additional major clades present in the tree (*n_uMCs*). Other important contributors included the proportion of sequences assigned to the donor major clade (*p_donorMC*), the patristic distance between the query and the closest sequence from an additional clade (*d*), and the proportion of sequences belonging to the recipient major clade (*p_recipientMC*). In contrast, the maximum topological distance within the tree (*max_topo*) and the mean proportion of additional major clades (*mean_p_uMCs*) showed comparatively lower contributions to model predictions.

Analysis of class-specific contributions revealed that distance-based features (*topo_d* and *d*) were particularly influential for distinguishing Multiple MC and Patchy phylogeny patterns. In addition, features describing the proportional distribution of taxa among major clades also played an important role in the classification process. Specifically, the proportions of sequences assigned to the recipient (*p_recipientMC*) and donor (*p_donorMC*) major clades showed strong contributions, primarily influencing the identification of Limited donor evidence and NoHGT patterns. Similarly, the feature *n_uMCs* contributed substantially to distinguishing iHGT patterns from the other evolutionary scenarios considered in this study.

Finally, other features, including the maximum topological distance (*max_topo*) and the mean proportion of additional major clades (*mean_p_uMCs*), exhibited moderate but consistent contributions across multiple classes.

Taken together, these analyses indicate that the Random Forest classifier achieves high discriminative performance across the evaluated phylogenetic patterns while relying on biologically interpretable signals derived from tree structure. The consistently high AUC values across all classes demonstrate the model’s ability to accurately distinguish among evolutionary scenarios. Both permutation importance and SHAP analyses further reveal that this predictive capacity is primarily driven by features describing phylogenetic topology and clade structure—particularly metrics such as *topo_d, d* and the number of additional major clades (*n_uMCs*). In addition, proportional descriptors of clade composition contribute complementary information that helps refine the classification of more complex phylogenetic patterns. Together, these results indicate that the extracted topological and clade-distribution features capture key evolutionary signals that enable robust differentiation among the phylogenetic patterns considered in this study.

### Performance with simulated data and unseen biological data

The features from phylogenetic trees derived from both the simulated datasets and the unseen real biological candidates were extracted using the feature-extraction pipeline and subsequently used as input for the trained Random Forest classification model. Under the controlled conditions of the simulated dataset, the model achieved a low misclassification rate of 7.8 % (detailed results are provided in Supplementary Table 10). When applied to the unseen real biological dataset, the model exhibited a slightly higher error rate of 10.43 %, reflecting the increased complexity and variability inherent to empirical data (detailed results are provided in Supplementary Table 11).

Analysis of the misclassified cases revealed that most errors involved confusion between the Multiple MC and Patchy phylogeny patterns. In the simulated dataset, approximately 20% of Multiple MC candidates were incorrectly classified as Patchy phylogeny. A similar trend was observed in the real biological dataset, where approximately 46% of Patchy phylogeny candidates were classified as Multiple MC. This pattern suggests that distinguishing between these two phylogenetic scenarios represents the most challenging task for the model, likely due to similarities in their topological and taxonomic signal.

To further evaluate the performance of the proposed classifier, the RF model was benchmarked against the AVP tool using two independent datasets: the simulated dataset used as a reference under controlled conditions and an unseen real biological dataset (Figure 4). The detailed results obtained with AVP are provided in Supplementary Table 12.

**Figure 4.**
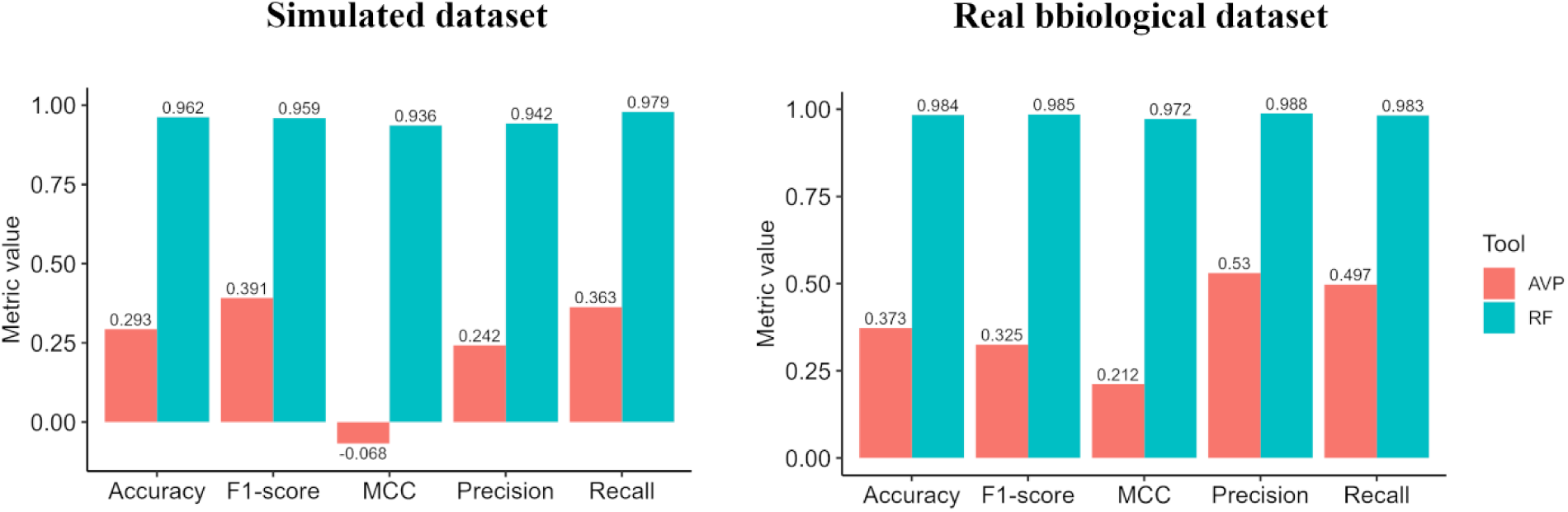
Comparison of classification performance between the Random Forest (RF) model and the AVP tool using simulated and real biological datasets. Performance was evaluated using accuracy, F1-score, Matthews correlation coefficient (MCC), precision, and recall. Bars represent the metric values obtained for each method. The dataset used for this comparison is provided in Supplementary Table 13.

In the simulated dataset, the RF model exhibited consistently high performance across all evaluated metrics, achieving an accuracy of 0.962, an F1-score of 0.959, a Matthews correlation coefficient (MCC) of 0.936, a precision of 0.942, and a recall of 0.979. In contrast, AVP showed substantially lower performance, with an accuracy of 0.293, an F1-score of 0.391, a precision of 0.242, and a recall of 0.363. Notably, AVP yielded a negative MCC value (−0.068), indicating poor agreement with the reference classifications and suggesting performance worse than random classification. These results indicate that the RF classifier accurately reproduced the expected classifications under controlled simulation conditions, whereas AVP showed limited predictive capability.

A similar pattern was observed for the real biological dataset. The RF model maintained very high performance, achieving an accuracy of 0.984, an F1-score of 0.985, an MCC of 0.972, a precision of 0.988, and a recall of 0.983. By contrast, AVP again showed markedly lower performance, with an accuracy of 0.373, an F1-score of 0.325, and an MCC of 0.212, while moderate values were observed for precision (0.530) and recall (0.497).

These results demonstrate that the RF-based approach consistently and substantially outperforms AVP across both simulated and real biological datasets, providing more accurate and reliable classification of phylogenetic patterns associated with iHGT.

## Discussion

The identification of interkingdom horizontal gene transfer (iHGT) has traditionally relied on manual inspection of phylogenetic trees, a process that is inherently subjective, difficult to reproduce, and challenging to scale to large datasets. As genomic data continues to expand, there is a growing need for approaches that can formalize expert-driven interpretation into reproducible and quantitative frameworks. In this study, we address this challenge by introducing a computational framework that translates recurrent phylogenetic patterns into a supervised classification problem. By combining biologically informed feature extraction with machine-learning approaches, we provide a scalable and interpretable method for classifying evolutionary scenarios associated with iHGT.

A central contribution of this work is the formalization of phylogenetic tree interpretation into a set of discrete, biologically meaningful patterns. Rather than restricting the analysis to a binary distinction between HGT and non-HGT, the proposed framework distinguishes among five recurrent evolutionary scenarios, including cases where the phylogenetic signal is insufficient to support definitive conclusions. This multi-class perspective more accurately reflects the complexity of evolutionary processes, where alternative explanations such as differential gene loss, incomplete lineage sorting, or multiple transfer events may generate similar topological signals (7,33). By explicitly incorporating these ambiguous cases into an Inconclusive category (Multiple MC and Patchy phylogeny), the framework avoids overinterpretation and provides a more nuanced representation of phylogenetic evidence.

The results demonstrate that phylogenetic tree topology and taxonomic composition can be effectively translated into quantitative descriptors that capture key evolutionary signals. Among the evaluated features, the topological distance between the query sequence and the closest sequence from an additional major clade (*topo_d*) emerged as the most influential predictor. This finding is biologically intuitive, as *topo_d* reflects the structural proximity of the query to lineages outside the inferred donor–recipient relationship. Small topological distances indicate that the query sequence clusters closely with taxa from additional major clades, a pattern consistent with the Multiple MC scenario, in which homologues from several lineages form a monophyletic group. Such configurations are often better explained by vertical inheritance followed by differential gene loss, which can mimic signals typically attributed to iHGT (7).

In contrast, larger topological distances indicate that sequences from additional major clades are more distantly positioned relative to the query and are distributed across different regions of the tree. This pattern is characteristic of Patchy phylogenies, where discontinuous taxonomic distributions may arise from a combination of evolutionary processes, including multiple independent horizontal transfer events, differential gene loss, or incomplete lineage sorting (34–37). Together, these results highlight that *topo_d* captures a key structural property of phylogenetic trees that is central to distinguishing between alternative evolutionary scenarios.

Similarly, the number of additional major clades (*n_aMCs*) provides a measure of lineage diversity within the tree, capturing the extent to which the phylogeny deviates from a simple two-clade structure. This feature plays a key role in the classification process, as it directly informs whether more complex evolutionary scenarios should be considered. In cases where no additional major clades are present (*n_aMCs* = 0), the phylogeny is restricted to a two-clade configuration, effectively excluding Inconclusive patterns such as Multiple MC and Patchy phylogeny. Under these conditions, the model can focus on distinguishing between iHGT and NoHGT, relying on complementary features such as the relative proportions of donor and recipient sequences.

Conversely, the presence of one or more additional major clades increases the topological and taxonomic complexity of the tree, expanding the range of plausible evolutionary scenarios. This information is therefore critical for guiding the classification toward Inconclusive patterns when appropriate. These findings are consistent with previous studies demonstrating that lineage diversity and topological structure provide informative signals for distinguishing among alternative evolutionary histories in phylogenetic trees (Kendall & Colijn, 2016; Xue et al., 2020).

Proportional features such as *p_donorMC* and *p_recipientMC* further contribute to the classification by reflecting the relative representation of taxa across donor and recipient major clades. These proportions are commonly considered during inferring horizontal gene transfer events, where the balance between donor and recipient sequences is used to assess the strength and direction of potential transfer events (20,22,38). iHGT scenarios are typically characterized by a well-represented donor clade relative to the recipient, whereas Limited donor evidence patterns involve a sparse representation of donor sequences, which reduces confidence in inferring the direction of transfer (39,40). Tree-based models such as Random Forest are well suited to capture these distinctions, as they partition the feature space into decision regions that implicitly reflect threshold-like relationships among variables (41,42). In this context, the model can learn combinations of proportional features that approximate the criteria used during expert-driven interpretation, without requiring the explicit definition of fixed thresholds.

These features operate in a complementary manner: *topo_d* captures local topological relationships, *n_aMCs* reflects global lineage diversity, and proportional descriptors quantify clade representation. Their combined contribution enables the model to reconstruct complex phylogenetic reasoning processes and to distinguish among evolutionary scenarios with high accuracy while maintaining biological interpretability.

The importance of these features is further supported by the permutation importance and SHAP analyses (Figures 2 and 3). Both approaches consistently identify *topo_d* as the dominant predictor, highlighting the central role of topological relationships in the classification process. Features such as *n_aMCs*, *p_donorMC*, and *p_recipientMC* also show substantial contributions, confirming that both lineage diversity and clade representation provide complementary information. Notably, SHAP analysis reveals that distance-based features are particularly influential in distinguishing complex patterns such as Multiple MC and Patchy phylogeny, whereas proportional features contribute more strongly to the classification of iHGT and Limited donor evidence scenarios. This concordance between model performance and feature interpretability supports the conclusion that the random forest (RF) classifier is not relying on spurious correlations but instead captures biologically meaningful signals derived from phylogenetic structure.

Tree-based ensemble methods, RF and XGBoost, consistently achieved the highest predictive performance. This result likely reflects their ability to model complex, non-linear interactions among features derived from phylogenetic tree topology and taxonomic composition (41,42). These descriptors are inherently dependent, as they capture different aspects of the same underlying structure, making them poorly suited to methods that assume linearity or feature independence.

In contrast, models such as Logistic Regression and Gaussian Naïve Bayes rely on assumptions that are unlikely to hold for phylogenetically derived features, while distance-based and kernel-based approaches may be sensitive to feature scaling and parameter selection (43–47). Neural network approaches may also be limited in this context due to the relatively modest dataset size, which can constrain their ability to learn stable and generalizable representations (48). These factors likely contributed to their comparatively lower performance.

Although RF and XGBoost showed similar predictive accuracy, RF was selected due to its greater interpretability. The independent construction of trees in RF facilitates stable and transparent estimates of feature importance, which is essential for linking model predictions to biological interpretation. In contrast, the sequential nature of XGBoost introduces more complex dependencies that are less straightforward to interpret at a global level (41,49,50).

Random Forest models are known for their stability and ability to effectively capture underlying patterns in complex datasets, as demonstrated in previous studies (51,52). In the present research, hyperparameter optimization did not lead to significant improvements in predictive performance, indicating that the model reached a performance plateau. This behavior suggests that the extracted features encode a strong and consistent phylogenetic signal that can be reliably learned without extensive parameter tuning, thereby reinforcing the robustness of the proposed framework.

The benchmarking analysis showed that the proposed RF classifier outperformed AVP in predicting iHGT events. This difference in performance may be explained by fundamental methodological distinctions between the two approaches. AVP relies on a two-step strategy based on implicit phylogenetic signals, initially using homology search results to compute metrics such as the Alien Index, HGT Index, Aggregate Hit Support (AHS), and outgroup percentage. While these measures provide useful heuristics, they are ultimately derived from sequence similarity comparisons between self (recipient lineage) and non-self (external lineage) hits, without explicitly incorporating phylogenetic structure (28,53–55).

Several studies have highlighted the limitations of such similarity-based approaches. For instance, processes such as amelioration can obscure ancient horizontal gene transfer events by homogenizing sequence composition over time, reducing detectable differences between donor and recipient lineages (56). Conversely, high sequence similarity does not necessarily reflect true evolutionary relatedness, as it may arise from convergent evolution or uneven taxonomic sampling (21,57,58).

Once candidates pass the first filter stage, the second AVP step is to build a phylogenetic tree using the homologous sequences. In this stage, not only the taxonomic annotation of leaf nodes is important, but also the phylogenetic position of these sequences relative to the iHGT candidate. AVP focuses on two branches—the sister branch of the gene of interest and the ancestral sister branch—and attaches donor-receptor tags only to those branches (28). While this simplifies decision-making and reduces computational load, restricting the analysis to these two branches may obscure the full evolutionary signal present across the rest of the tree. Ignoring remote branches and broader topological patterns could mask other phylogenetic scenarios, including incomplete lineage sorting, hybridization or gene loss that would otherwise support alternative evolutionary hypotheses or challenge iHGT inferences (7,34,59).

Rather than focusing only on local branches, the proposed RF classifier incorporates information from the entire phylogenetic tree during feature extraction. This global perspective enables the systematic quantification of key structural and taxonomic properties, including topological distances, the number of major clades, and the relative distribution of taxa across lineages. By capturing signals from the full tree rather than a restricted topological neighborhood, the model can represent a broader spectrum of evolutionary patterns.

This comprehensive integration of phylogenetic information reduces the risk of misclassifying scenarios that superficially resemble iHGT but may instead arise from alternative evolutive processes. Consequently, the proposed framework provides a more robust and nuanced interpretation of evolutionary histories, improving discrimination between true iHGT events and other phylogenetic configurations that can generate similar local signals.

Over the years, the detection of horizontal gene acquisition in eukaryotes has been extensively discussed in the literature, giving rise to a variety of tools that adopt distinct approaches—parametric, implicit and explicit phylogenetic methods—for iHGT inference (20,56,60).

According to Richards et al. (2011), the evaluation of discordance between gene tree and species tree remains the gold standard for detecting iHGT candidates. However, a high error rate in inferring iHGT events due to gene-tree reconciliation artifacts has since been documented (32). Furthermore, database bias, taxon sampling, and methodological decisions all contribute to uncertainty in iHGT inference, as demonstrated by Aguirre-Carvajal & Armijos-Jaramillo (2026). Together, these factors highlight the lack of a general consensus on the criteria required to confidently identify a given scenario as iHGT.

The application of artificial intelligence represents a promising direction for advancing the understanding of horizontal gene transfer in eukaryotes. Although recent studies have demonstrated the potential of these approaches to yield meaningful insights, to date no method has explicitly leveraged machine learning to infer iHGT candidates within a fully phylogenetic framework (62). In this context the proposed tool represents a novel and promising approach for distinguishing between iHGT and NoHGT candidates. However, as shown in Figure 1 and the benchmarking results, the model struggles to differentiate between Multiple MC and Patchy phylogeny patterns, which are inherently challenging to classify. The proposed random forest classifier may be improved by expanding the feature set or exploring alternative feature combinations, which could enhance its performance and discriminatory power (63).

Finally, an important contribution of this study is the development of a validation resource for benchmarking iHGT detection methods. To date, there is no widely accepted reference dataset for robust evaluation in this domain, as noted by Wijaya et al. (2025). To address this gap, we generated a simulated dataset of phylogenetic trees representing iHGT-related scenarios using HgtSIM. Beyond its utility for benchmarking, this dataset provides insights into the biological plausibility of different phylogenetic patterns.

The simulations indicate that certain topological configurations commonly interpreted as evidence of iHGT may require a series of highly specific and biologically unlikely events as was already highlighted in Bremer et al. (2025) study. For instance, reproducing the Multiple MC pattern required simulating independent gene transfer events from bacteria into multiple eukaryotic major clades. Biologically, this would imply that the same bacterial gene was transferred into several distinct eukaryotic lineages at the same time, a scenario that is possible but likely rare. Similarly, generating Patchy phylogenies required multiple independent transfer events involving different bacterial donors and distinct eukaryotic recipients, implying concurrent or sequential transfers across diverse lineages in the same period of time. These simulations suggest that interpreting such complex topologies as direct evidence of iHGT may overestimate the likelihood of these events.

This study demonstrates that phylogenetic tree interpretation can be translated into a consistent and data-driven process, enabling the systematic evaluation of complex evolutionary scenarios.

By capturing topological structure, lineage diversity, and taxonomic representation within a unified framework, the approach provides a more rigorous basis for distinguishing between true iHGT signals and alternative evolutionary processes that can generate similar patterns.

Importantly, the results highlight that some phylogenetic configurations commonly interpreted as evidence of iHGT may instead reflect alternative evolution scenarios, underscoring the need for cautious and context-aware inference. In this sense, the proposed method not only improves predictive performance but also contributes to refining the criteria by which iHGT events are evaluated.

Despite these advantages, the proposed tool has its own inherent limitations, as it is built on our current knowledge, experience, and evaluation criteria; if these criteria change, the model’s accuracy is expected to decline. Future developments should focus on expanding the diversity of training data, incorporating additional sources of evolutionary evidence, and further exploring ambiguous phylogenetic patterns that remain difficult to resolve. Advancing in these directions will be essential for establishing more robust and widely applicable standards for iHGT detection in the era of large-scale comparative genomics.

## Methodology

### Conceptual framework for phylogenetic pattern classification

Manual inspection of phylogenetic trees has long been a central approach for identifying interkingdom horizontal gene transfer (iHGT) events. During the examination of large collections of atypical gene trees across multiple studies investigating iHGT, recurrent topological configurations were consistently observed. These configurations correspond to distinct evolutionary scenarios identifiable through tree topology. Based on these observations, five recurrent phylogenetic patterns were identified: iHGT, Limited donor evidence, Multiple major clades (MC), Patchy phylogeny, and NoHGT (39).

To facilitate the interpretation of phylogenetic tree topology and the identification of recurrent phylogenetic patterns, taxa present in each tree were grouped into broader taxonomical categories referred to as major clades (MCs) (Table 2). In total, organisms were classified into eight MCs, each encompassing multiple associated minor clades. These groupings are based on broad taxonomic categories derived from the NCBI Taxonomy database and previous studies (2,6). This hierarchical classification provides a standardized framework for summarizing gene tree taxonomic composition and facilitating comparisons among distantly related lineages.

**Table 2.**
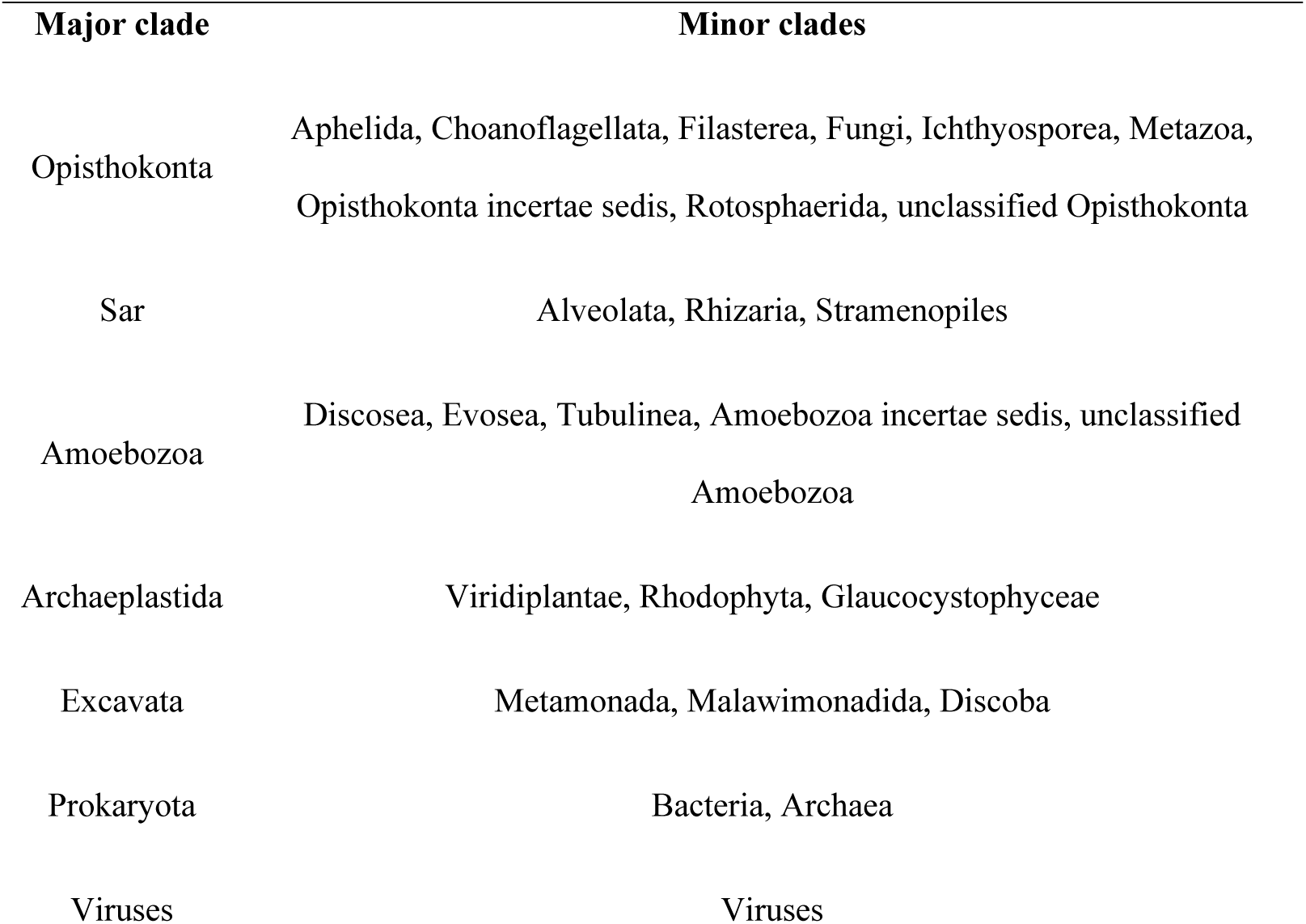
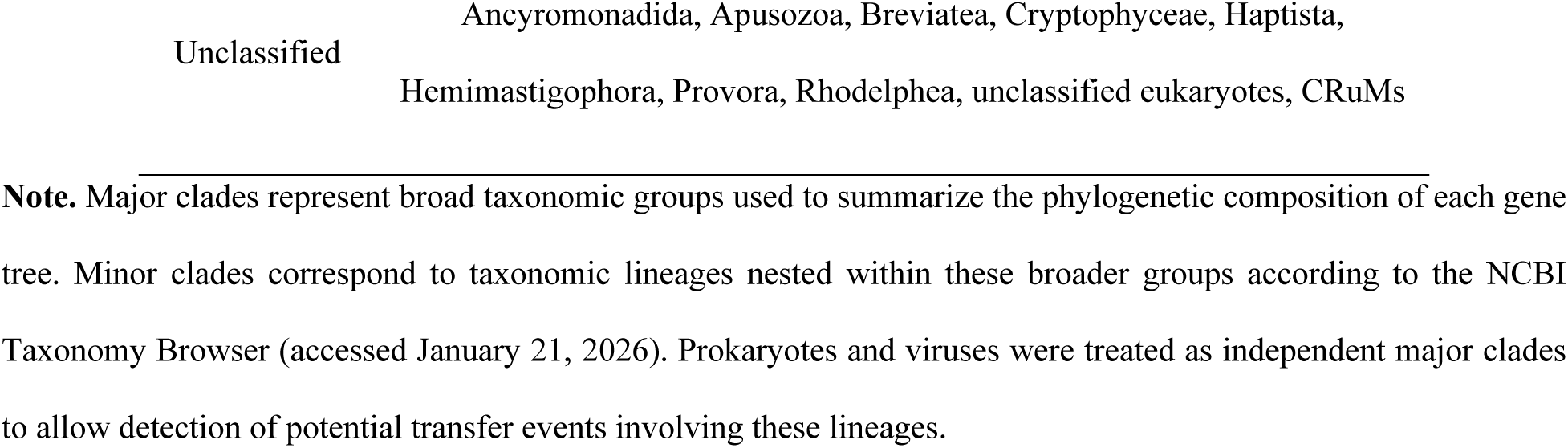
Classification scheme of major clades and associated minor clades applied in this study.

Among the MCs, six correspond to major eukaryotic lineages. One of these categories (*Unclassified*) groups eukaryotic taxa that cannot be confidently assigned to a defined clade. In addition, prokaryotes and viruses were treated as independent MCs to allow detection of transfer scenarios involving these lineages when present in the phylogeny.

Phylogenetic patterns were defined based on gene tree topology and the distribution of sequences across MCs, with each pattern representing a distinct evolutionary scenario. To illustrate iHGT, NoHGT, Limited donor evidence, Multiple MC and Patchy phylogeny patterns, conceptual phylogenetic topologies are shown in Figure 5.

**Figure 5.**
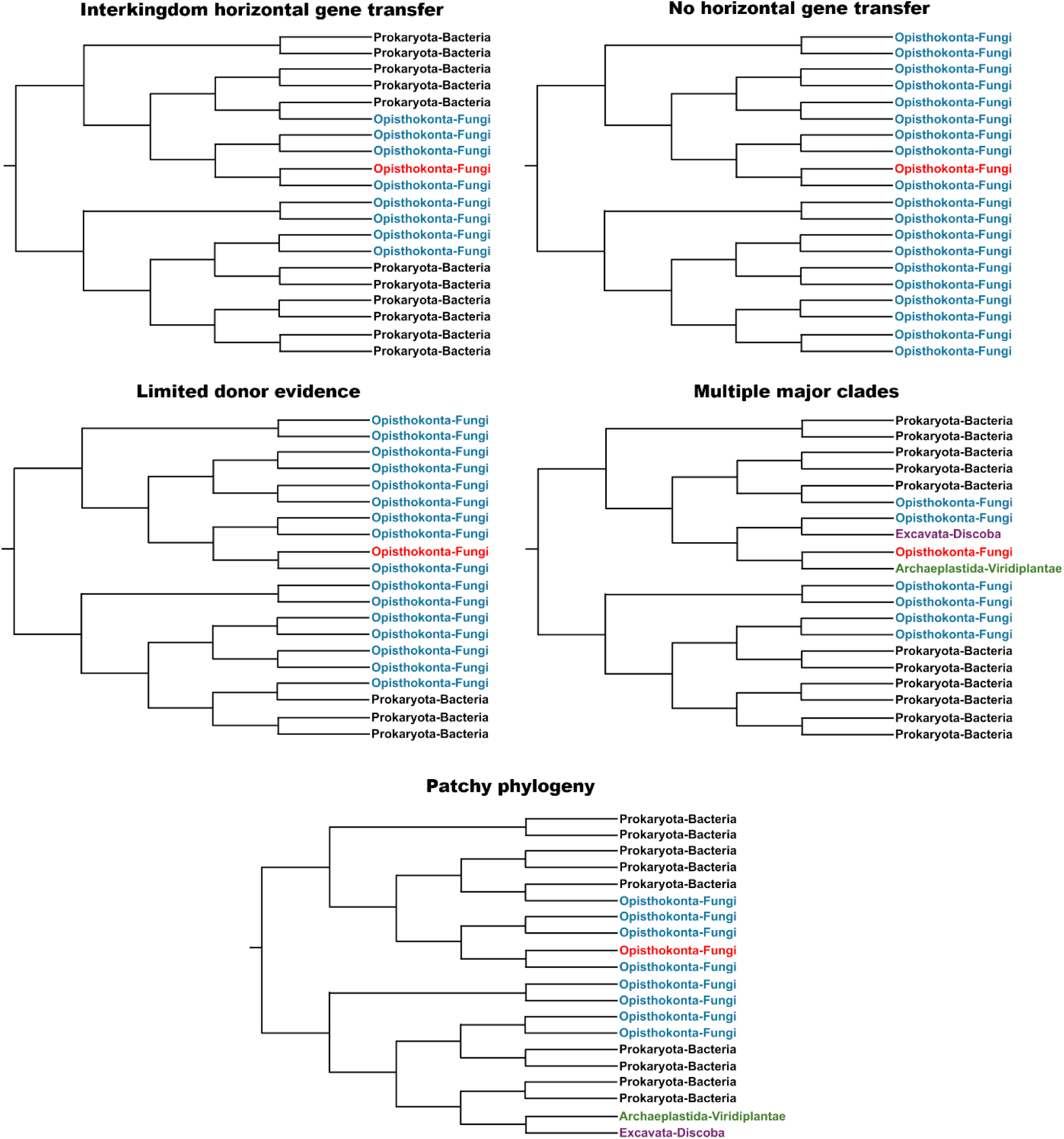
Schematic gene trees examples illustrate five recurrent evolutionary scenarios: interkingdom horizontal gene transfer (iHGT), No horizonal gene transfer (NoHGT), Limited donor evidence, Multiple major clades (Multiple MC), and Patchy phylogeny. Trees are based on a representative example of gene transfer from bacteria (Prokaryota) to fungi (Opisthokonta). Colored labels indicate major clades, and the query sequence is highlighted in red. These conceptual topologies are intended to illustrate the distinguishing features of each phylogenetic pattern.

The iHGT scenario represents gene flow occurring exclusively between two major clades. This pattern is characterized by phylogenetic trees containing sequences from only two MCs: a putative donor and a recipient. In such trees, the query sequence belongs to the recipient MC and forms a monophyletic group nested within, or positioned as sister to, sequences from the donor MC. This topological configuration is consistent with the acquisition of the gene through an iHGT event and difficult to explain by alternative hypothesis.

The NoHGT class represents phylogenetic trees in which sequences belong to a single MC. In these cases, the tree contains query homologs sequences exclusively from that clade, with no representatives from other MCs. Because no cross-clade phylogenetic relationships are observed, the topology does not provide evidence supporting gene acquisition from another MC. This configuration is therefore interpreted as consistent with vertical inheritance within the lineage rather than iHGT. Accordingly, these cases are classified as NoHGT.

Limited donor evidence, Multiple MC, and Patchy phylogeny correspond to cases in which the phylogenetic signal is insufficient to confidently determine whether a gene candidate originated through iHGT. In the Limited donor evidence pattern, phylogenetic trees contain sequences from two MCs, but the putative donor clade is represented by relatively few sequences compared with the recipient clade. This limited representation may indicate that the putative donor lineage is underrepresented in current databases, or that the direction of gene flow differs from the initial hypothesis. Because the scarcity of donor sequences reduces phylogenetic resolution and weakens confidence in inferring the direction of transfer, these cases do not provide sufficient evidence to robustly support an iHGT scenario.

The Multiple MC and Patchy phylogeny patterns correspond to phylogenetic trees containing sequences from three or more MCs. In the Multiple MC pattern, the query sequence clusters within a monophyletic group including sequences from the donor, recipient, and additional major clades. This topology is consistent with vertical inheritance followed by differential gene loss across lineages. Alternatively, an iHGT hypothesis would require invoking an ancient transfer followed by multiple independent gene losses, making the evolutionary scenario difficult to resolve.

In contrast, the Patchy phylogeny pattern is characterized by additional major clades distributed across different parts of the tree rather than forming a monophyletic group with the query sequence. Such topological configurations may result from several evolutionary processes, including differential gene loss, incomplete lineage sorting, or multiple independent horizontal gene transfer events between kingdoms.

Because gene tree topology alone cannot reliably distinguish among these alternative evolutionary scenarios, candidates classified as Limited donor evidence, Multiple MC, or Patchy phylogeny were considered Inconclusive.

### Feature extraction from phylogenetic trees

The tool was designed to implement a supervised classification model capable of emulating the manual analysis performed by researchers when examining gene tree topologies and assigning them to distinct phylogenetic patterns. Studies investigating iHGT have shown that gene tree interpretation primarily relies on two core properties. The first is the taxonomic composition of the sequence homologues (6,22,23). The second is the phylogenetic relationships among these sequences, particularly their position relative to the query sequence and whether they form a monophyletic group or instead appear as dispersed lineages that open different options difficult to determine with a single phylogeny (28).

To translate the criteria used during manual phylogenetic inspection into a machine-learning framework, a set of quantitative features were extracted from gene trees. These features capture both the taxonomic composition of the sequences and their phylogenetic relationships. The descriptors were specifically designed to formalize expert-driven interpretation of gene tree topologies into a quantitative framework. While their conceptual basis is grounded in previously described criteria for identifying iHGT events, their formal definition and integration into a machine-learning classification pipeline represent a novel contribution of this study.

For this purpose, sequences were classified according to their affiliation with the donor and recipient MCs. Any MC that differs from both the putative donor and the query recipient MC are hereafter referred as additional MC. An illustrative representation of the phylogenetic features extracted from a gene tree is shown in Figure 6.

**Figure 6.**
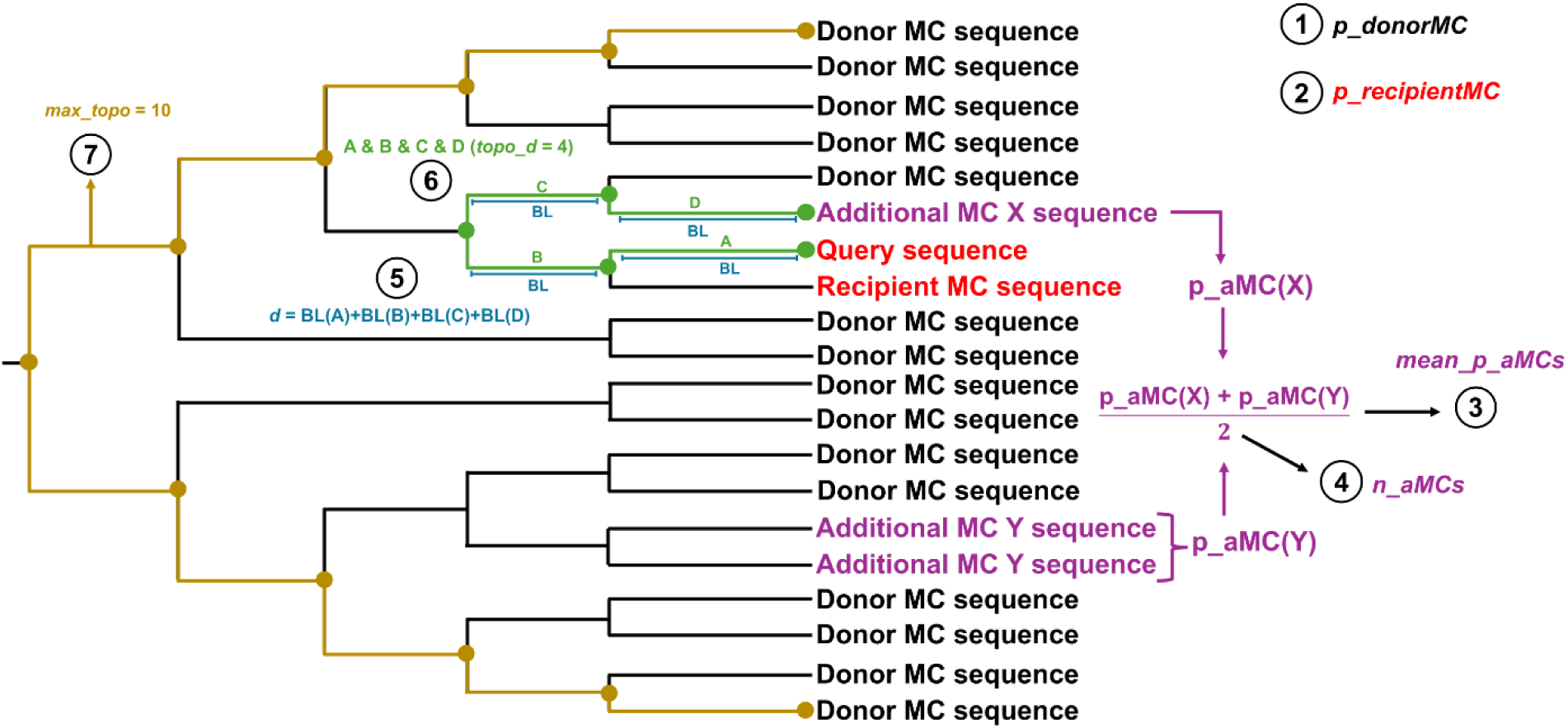
The schematic tree represents a hypothetical phylogeny containing sequences from the inferred donor major clade, recipient major clade, and additional major clades. The candidate (query) sequence and recipient major clade (MC) sequences are highlighted in red, whereas sequences belonging to the donor, recipient, and additional clades are indicated in black, and purple, respectively. Numbered annotations represent the quantitative features extracted from the phylogenetic tree. (1) Proportion (p) of sequences belonging to the donor MC (*p_donorMC*). (2) Proportion of sequences belonging to the recipient MC (*p_recipientMC*). (3) Mean proportion of sequences belonging to additional (a) major clades (MCs) (mean_p_aMCs), calculated by first determining the proportion of terminal nodes for each additional MC (each distinct taxonomic group different from the donor and recipient MCs) and then averaging these proportions across all additional MCs present in the tree. (4) Number of additional major clades detected in the tree (*n_aMCs*). (5) Patristic distance (*d*) and (6) topological distance (*topo_d*) between the query sequence and the closest sequence from an additional major clade. Patristic distance corresponds to the sum of branch lengths (BL) along the path connecting two terminal nodes in the tree. In contrast, topological distance represents the number of edges (branches) separating the same pair of nodes, independent of branch length. (7) Maximum topological distance (*max_topo*) within the phylogeny, representing the structural diameter of the tree. Together, these features capture both the taxonomic composition and the phylogenetic relationships that underpin the manual interpretation of gene tree topologies. Branch lengths are not drawn to scale, and the topology is shown for illustrative purposes.

The first group of features describes the taxonomic composition of each gene tree. These variables quantify the relative representation of sequences belonging to the donor, recipient and additional MCs. Together, they summarize the distribution of lineage diversity within the tree, which constitutes a key criterion used during manual classification of phylogenetic patterns.

Specifically, four features were computed to capture these aspects of taxonomic composition. The proportion of the major donor clade (*p_donorMC*; 1 in Figure 6) represents the fraction of terminal nodes assigned to the putative donor major clade. The proportion of the major recipient clade (*p_recipientMC*; 2 in Figure 6) corresponds to the fraction of taxa belonging to the recipient clade. To account for additional lineage diversity, the average proportion of additional major clades (*mean_p_aMCs*; 3 in Figure 6) was calculated. First, the proportion of sequences belonging to each distinct additional MC was computed. These values were then averaged across all additional MCs present in the phylogeny. Finally, the number of additional major clades (*n_aMCs*; 4 in Figure 6) represents the total count of distinct additional clades detected in the tree.

The second group of features quantifies the phylogenetic relationships between the query sequence and sequences belonging to additional MCs in the tree. During manual phylogenetic inspection, researchers evaluate how closely the query sequence clusters with sequences from such clades, as these relationships can influence the inferred evolutionary scenario. To capture this information computationally, two complementary distance metrics were calculated. Both metrics were normalized by the corresponding tree diameter to ensure comparability across trees with different depths and branch-length scales.

The first metric consists of patristic distances (*d*; 5 in Figure 6), which measure evolutionary divergence as the sum of branch lengths separating two taxa. The second metric consists of topological distances (*topo_d*; 6 in Figure 6), which quantify structural separation as the number of edges connecting two taxa in the tree. Together, these distance-based features quantify the phylogenetic proximity between the query sequence and lineages belonging to additional MCs. The presence and relative placement of such sequences may indicate broader lineage distributions that complicate interpretations based on simple donor–recipient transfer patterns. Additionally, the maximum topological distance within the tree (*max_topo*; 7 in Figure 6) was extracted as a separate feature to characterize the overall structural extent of the phylogeny.

All features included in the model were derived directly from phylogenetic trees using a fully reproducible pipeline implemented in Python v3.10.12 (Figure 7). The pipeline accepts trees in either NEXUS or Newick format, and branch lengths are expressed in substitutions per site. Phylogenies were parsed using the *Phylo* module of Biopython v1.85 (64).

**Figure 7.**
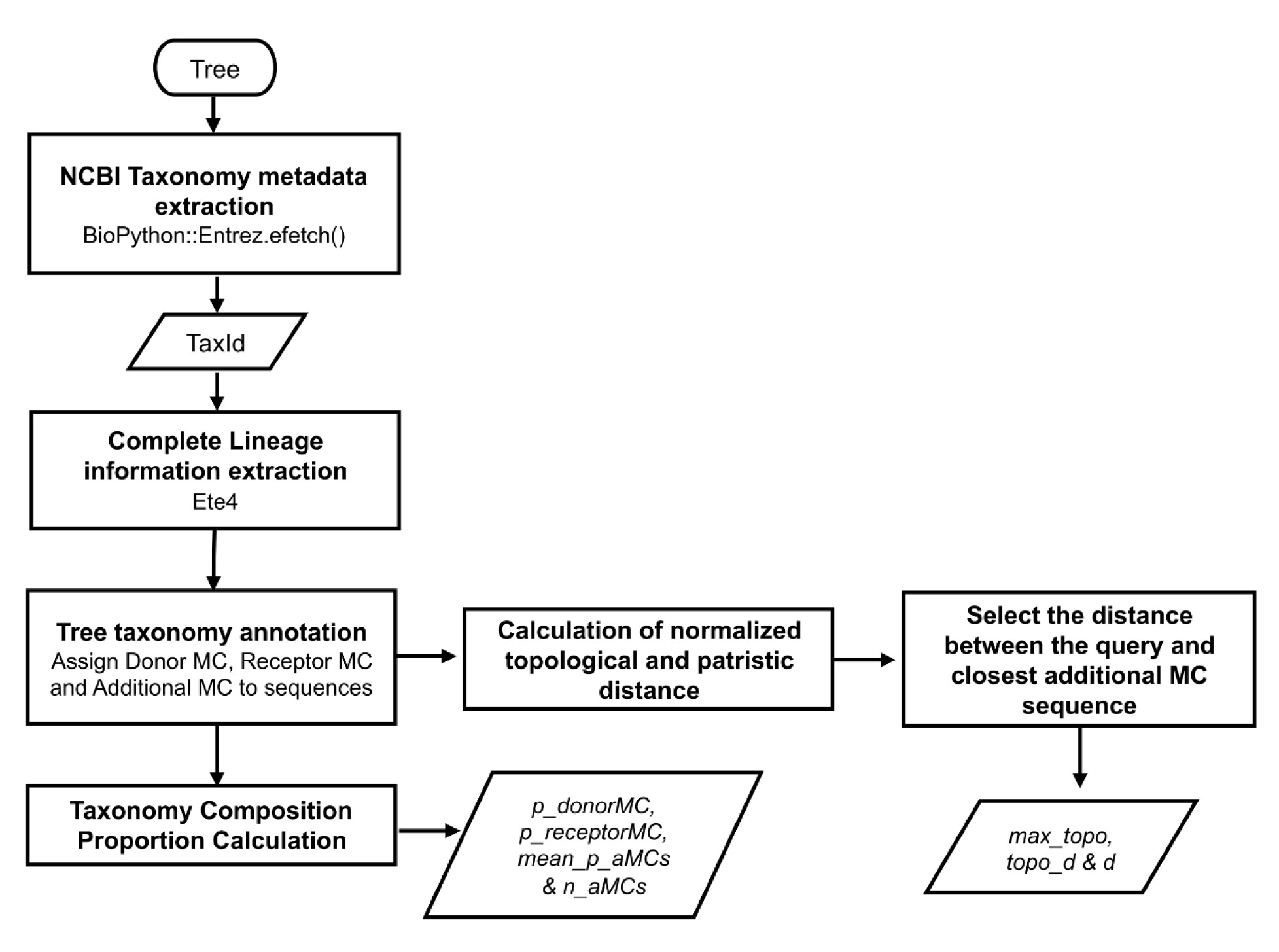
Overview of the workflow used for taxonomic annotation and extraction of phylogenetic features. The workflow begins with a phylogenetic tree, from which NCBI Taxonomy metadata are retrieved from NCBI sequence identifiers to obtain taxonomic identifiers. Complete lineage information is then extracted using ETE4, enabling detailed taxonomic annotation of each sequence. Based on this information, each sequence is assigned to the donor, recipient, or an additional major clade (MC). Next, normalized patristic and topological distances are computed between the query sequence and all other sequences in the tree. From these values, the minimum distance between the query and sequences from additional MCs is retained, yielding the features *d* and *topo_d*. The maximum topological distance within the tree (*max_topo*) is also calculated. Finally, taxonomic composition features are computed, including the proportions of sequences belonging to the donor MC and recipient MC (*p_donorMC*, *p_recipientMC*), as well as the mean proportion and number of additional MCs (*mean_p_uMCs* and *n_uMCs*). Together, these variables constitute the feature set used for downstream classification of phylogenetic patterns.

Each terminal node corresponds to an NCBI protein accession, enabling retrieval of associated taxonomic metadata. For each accession, the corresponding GenBank record was obtained using the Entrez efetch utility implemented in Biopython. The NCBI taxonomy identifier (TaxID) was extracted from the record and used to retrieve the complete taxonomic lineage through the NCBITaxa module of the ETE Toolkit v4.0.0b2 (65).

Based on the retrieved taxonomic lineages, sequences were assigned to their corresponding major and minor clades according to the classification scheme summarized in Table 2. Following this classification, all terminal taxa in each tree were grouped into three analytical categories: (i) sequences belonging to the major recipient clade, (ii) sequences assigned to the putative major donor clade, and (iii) sequences affiliated with additional major clades.

To define the donor major clade (MC), the recipient MC—provided by the user—is first excluded from the set of MCs present in the tree, after which the most abundant remaining MC is selected as the donor MC. Following taxonomic annotation, patristic and topological distances were computed using the Phylo module of Biopython and normalized by the corresponding tree diameter. For each distance metric, the minimum distance between the query sequence and sequences from additional MCs was retained, yielding the features *d* and *topo_d*. Additionally, the maximum topological distance within the tree (*max_topo*) was extracted.

Finally, features describing the taxonomic composition of each tree were computed. Specifically, the proportions of sequences assigned to the donor (*p_donorMC*) and recipient (*p_recipientMC*) major clades were estimated. For additional MCs, individual proportions were computed and used to derive mean_p_uMCs and n_uMCs. All proportion calculations were restricted to sequences for which a complete taxonomic lineage could be successfully retrieved and assigned.

The resulting feature set was compiled into a tab-delimited output file representing the final output of the pipeline. The parameters used to run the pipeline are summarized in Supplementary Table 1.

### Supervised model training and performance evaluation

Model training was conducted using a manually curated phylogenetic trees publicly available in the repository associated with the study of (6). The dataset comprises 1,012 phylogenetic trees, each represented by a set of seven quantitative features extracted from gene tree topology and taxonomic composition. The model was trained to classify five phylogenetic patterns (iHGT, NoHGT, Limited donor evidence, Multiple MC, and Patchy phylogeny), which were later grouped into three broader categories (iHGT, NoHGT, and Inconclusive) for benchmarking comparisons. The distribution of phylogenetic patterns and their corresponding class assignments is summarized in Table 3. The dataset is accessible through Zenodo (https://doi.org/10.5281/zenodo.17552462).

**Table 3.** Distribution of phylogenetic patterns.

| **Grouped category** | **Phylogenetic pattern**<br>**(training class)** | **Count** |
| --- | --- | --- |
| iHGT | iHGT | 332 |
| NoHGT | NoHGT | 73 |
| Inconclusive | Limited donor<br>evidence | 117 |
|  | Multiple MC | 176 |
|  | Patchy phylogeny | 314 |

Although the dataset is not balanced, the class imbalance ratio—defined as the ratio between the most represented class (iHGT) and the least represented class (NoHGT) at the phylogenetic pattern level—is approximately 4.5:1, which remains below levels commonly regarded as severe class imbalance in machine learning (e.g., ratios exceeding 10:1 or even 100:1) (66,67). Under these conditions, the original class distribution was preserved and no artificial rebalancing procedures were applied, as moderate imbalance can often be handled effectively by ensemble methods (68).

After completing the feature-extraction pipeline, seven supervised learning algorithms were evaluated to identify the model best suited for classifying the phylogenetic patterns defined above. The models evaluated included Random Forest (RF), Extreme Gradient Boosting (XGBoost), Multilayer Perceptron (MLP), Logistic Regression (LR), Gaussian Naïve Bayes (GNB), K-Nearest Neighbors (KNN), and Support Vector Classifier (SVC). All models were implemented in Python using the scikit-learn library v1.7.2 and XGBoost package v3.2.0.

Model-specific parameter settings are summarized in Table 4. Unless otherwise specified, all remaining parameters were set to their default values as implemented in the corresponding software versions used in this study. Minor adjustments were applied when required to ensure reproducibility or compatibility with multi-class classification.

**Table 4.** Machine learning models evaluated for phylogenetic pattern classification, including implementation details and parameter settings.

| Model | Function | Parameters |
| --- | --- | --- |
| RF | RandomForestClassifier() | n_estimators=100, random_state=42, |
| XGBoost | XGBClassifier() | n_estimators=100, eval_metric="mlogloss", random_state=42 |
| MLP | MLPClassifier() | max_iter=5000, hidden_layer_sizes=(100,), random_state=42 |
| LR | LogisticRegression() | solver="lbfgs", max_iter=5000, random_state=42 |
| GNB | GaussianNB() | var_smoothing=1e-09 |
| KNN | KNeighborsClassifier() | n_neighbors=5 |
| SVC | SVC() | kernel='rbf', probability=True, random_state=42 |

**Note.** Machine-learning models evaluated during the training phase, including their Python implementations and key parameter settings. Parameters not explicitly listed were kept at their default values as implemented in the corresponding software versions. Reported settings include parameters relevant to model configuration and reproducibility.

All models were trained using a standardized machine-learning pipeline implemented with the *Pipeline* class in scikit-learn. The pipeline consisted of two steps: (1) missing-value imputation using *SimpleImputer* with a constant strategy (*fill_value* = –1), and (2) the classification model under evaluation. Missing values arose when no homologs from additional major clades were present in the tree. This procedure ensured consistent preprocessing across all models.

To obtain robust and statistically comparable estimates of model performance, we implemented a repeated stratified k-fold cross-validation scheme using the *RepeatedStratifiedKFold* function. Specifically, five folds (*n_splits* = 5) were used, resulting in an 80% training set and a 20% test set for each split. The procedure was repeated 10 times (*n_repeats* = 10) with different random partitions (*random_state* = 42), producing 50 performance estimates per model while preserving class proportions across splits.

For each repetition, model performance was evaluated on the corresponding test partition using five metrics: area under the receiver operating characteristic curve (AUC–ROC), precision, recall, F1-score, and accuracy. For the multi-class setting, AUC–ROC was calculated using a one-vs-rest strategy. AUC-ROC, precision, recall, and F1-score were computed using weighted averaging to account for class imbalance, ensuring that each class contributed proportionally according to its frequency in the dataset. Accuracy was calculated as the proportion of correctly classified instances. These metrics capture discrimination ability (AUC–ROC), class-balanced predictive performance (precision, recall, F1-score), and overall accuracy. The resulting performance estimates were subsequently used for statistical comparison among models.

The experimental setup followed a completely randomized repeated-measures design, in which each classification model (Table 4) was treated as an independent factor and performance metrics as dependent variables. Repeated measurements arise from the 50 performance estimates obtained through the repeated stratified cross-validation procedure, enabling statistical comparison of model performance across identical resampling partitions.

To statistically compare model performance across the repeated cross-validation evaluations, analyses were conducted in RStudio using non-parametric tests appropriate for repeated-measures designs, implemented with the R v4.5.3 packages stats v4.2.2 and PMCMRplus v1.9.12. Differences among classification models were first assessed using Friedman’s test. When significant differences were detected, pairwise comparisons were performed using the Nemenyi post hoc test. Bonferroni correction was applied to account for multiple comparisons. Statistical significance was assessed at α = 0.05.

The model exhibiting the best performance in the preliminary comparison and was therefore selected for hyperparameter optimization. Grid search and randomized search were performed using *GridSearchCV* and *RandomizedSearchCV* with the same repeated stratified cross-validation scheme described above. The optimized configurations were subsequently evaluated using the same statistical comparison framework applied in the initial model assessment.

Once the best-performing model was identified, a detailed model interpretability analysis was conducted. This included: (i) class-specific AUC-ROC curves to evaluate discriminatory performance for each phylogenetic category; (ii) permutation feature importance, computed with sklearn.inspection, to quantify the decrease in predictive performance associated with permuting each feature; and (iii) SHAP (SHapley Additive exPlanations) analysis, implemented using the shap v0.49.1 package, to characterize both global and instance-level feature contributions. Together, these analyses provide a comprehensive and interpretable assessment of the factors driving model decisions.

### Simulation of phylogenetic patterns

To validate model performance under controlled conditions, we simulated a total of 1,000 phylogenetic trees, comprising 200 replicates for each phylogenetic pattern described in Table 3. Simulations were conducted using HgtSIM v1.1.0 (69). To ensure consistency with the intended application scope, simulations were restricted to single-celled organisms, as the framework targets microbial systems. The set of organisms included in the simulations is provided in Supplementary Table 2.

The simulation framework was designed to reproduce the phylogenetic patterns under controlled evolutionary scenarios in which bacterial organisms, used here as representatives of Prokaryota, acted as donors and transfer genes into fungi (Opisthokonta). Ten genes widely conserved in bacteria—*murA, murB, murC, murD, murE, murF, lpxA, rpoD, fliC,* and *ddl* (70–74)—were selected as donor loci. These genes were selected as representative bacterial loci for simulating donor-derived sequences, rather than to imply exclusivity to prokaryotes.

To obtain donor sequences, a reference protein accession was first identified through keyword-based searches in the NCBI Protein database (accessed January 15, 2026), and a single representative protein sequence was selected to serve as the donor template in subsequent simulations. Protein sequences were used instead of directly retrieving nucleotide sequences to ensure consistent coding regions, as preliminary tests using nucleotide records resulted in reading frame inconsistencies during HgtSIM processing. The corresponding sequences were then retrieved using Entrez Direct v24.0 (efetch utility). As HgtSIM operates on nucleotide sequences, the selected amino acid sequences were back-translated into coding DNA sequences using EMBOSS v6.6.0.0 (backtranseq utility). In parallel, reference genome assemblies of the recipient organisms were obtained using the NCBI Datasets tool v18.14.0. This information was subsequently used to perform the HgtSIM simulations.

To facilitate the interpretation of the simulation design, a schematic overview of the workflow and the resulting phylogenetic patterns is provided in Figure 8. This figure illustrates how donor sequences of prokaryotic origin are introduced into recipient genomes under different evolutionary scenarios, as well as how homologous sequences are retrieved and used to reconstruct phylogenetic trees. Each simulated scenario corresponds to a distinct phylogenetic pattern defined by the distribution of homologues across MCs and the topology of the inferred gene trees.

**Figure 8.**
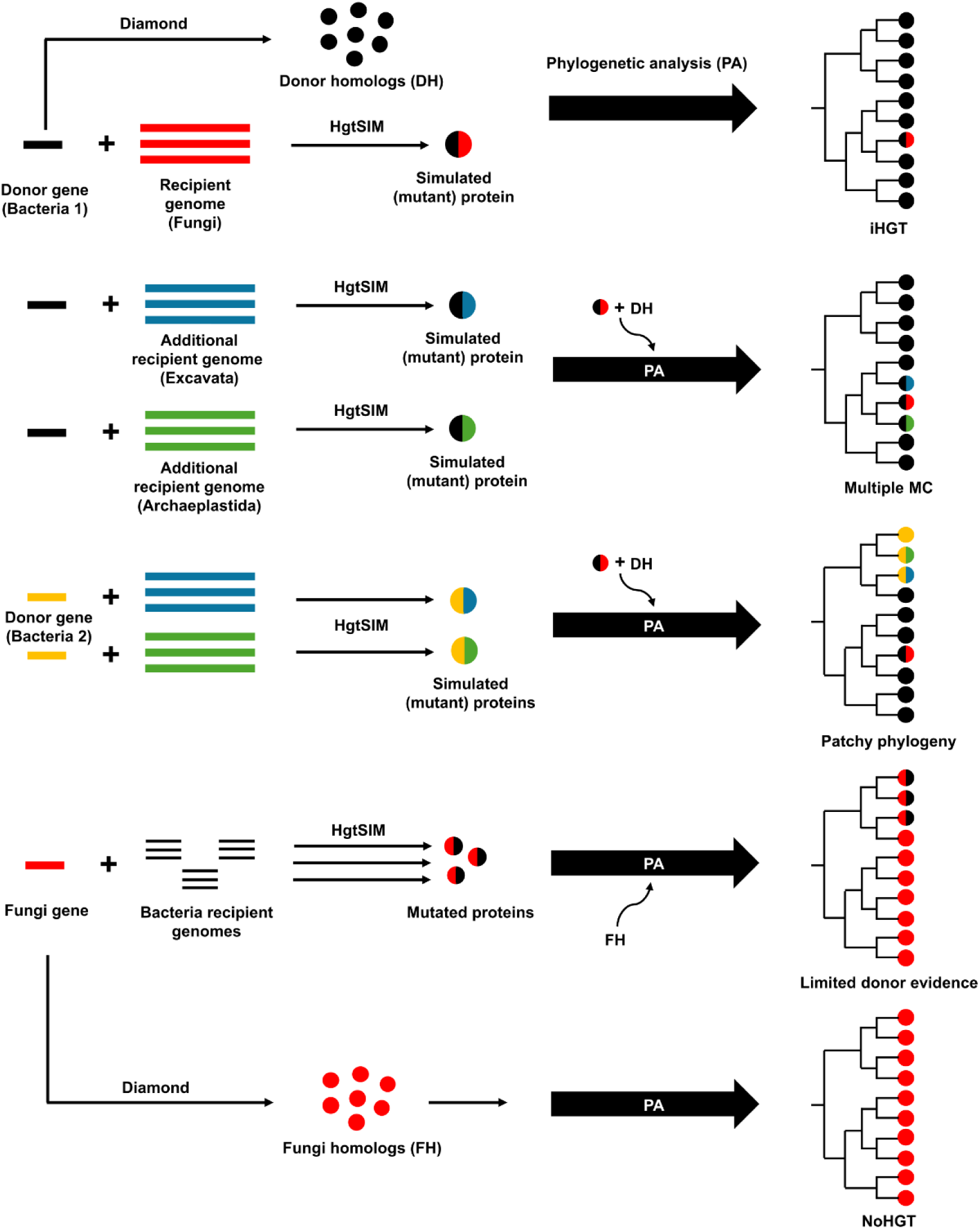
Schematic representation of how studied phylogenetic patterns were simulated using HgtSIM. Donor genes of prokaryotic origin (Bacteria 1 or Bacteria 2) are introduced into recipient fungal genomes (Opisthokonta) or other eukaryotic lineages to generate distinct evolutionary patterns. Homologous sequences are retrieved using DIAMOND, and phylogenetic analysis (PA) is performed to reconstruct gene trees. The resulting topologies correspond to five patterns: (i) interkingdom horizontal gene transfer (iHGT), where fungal sequences cluster within prokaryotic homologues; (ii) Multiple MC, where homologues are distributed across multiple major clades (MCs) but form a monophyletic group; (iii) Patchy phylogeny, characterized by discontinuous distribution of homologues across clades; (iv) Limited donor evidence, where donor sequences are sparsely represented; and (v) NoHGT, where no horizontal transfer signal is observed. Abbreviations: DH, donor homologs; FH, fungal homologs; PA, phylogenetic analysis.

As illustrated in Figure 8, these scenarios were implemented as follows. Donor sequences were horizontally introduced into 20 distinct fungal genomes, generating a total of 200 baseline simulated iHGT events. To generate the Multiple MC pattern, the transferred gene was subsequently introduced into taxa belonging to MCs distinct from Opisthokonta thus producing trees in which homologues are distributed across multiple major lineages while clustering in a monophyletic clade. Finally, to simulate the Patchy phylogeny pattern, the gene transferred into fungi originated from a single prokaryotic donor species, whereas homologues in other major clades were derived from different prokaryotic species independently carrying the same gene. This design generated discontinuous phylogenetic distributions consistent with patchy evolutionary histories.

To simulate the Limited Donor Evidence pattern, 200 fungal genes were selected and transferred into a set of prokaryotic organisms, thereby generating trees in which the donor clade is sparsely represented. To generate the NoHGT pattern, the same 200 fungal genes were retained without introducing any horizontal transfer event. To ensure that no transfer scenario was duplicated, each simulation was assigned a unique identifier. The structure of these simulations was generated using a custom R script. The complete set of simulation configurations and relevant metadata to run HgtSIM is provided in Supplementary Table 3.

After compiling all required inputs for each recipient genome, HgtSIM was executed using the default mutation ratio parameter and varying mutation intensity levels (-i). The parameter -t specifies the nucleotide sequence to be transferred in FASTA format, whereas -d defines a distribution file indicating the gene identifier to be transferred and the corresponding recipient genome identifiers. The directory path containing the recipient genome assemblies is provided using the -f option. The parameter -x specifies the file extension of the recipient genome assemblies. Detailed instructions for running HgtSIM are available in the project documentation (https://github.com/songweizhi/HgtSIM). Simulations were performed using the following command-line structure: HgtSIM -t [gene.fasta] -d [distribution.txt] -f [recipient_genome_directory] -r 1-0-1-1 -x fna -i [mutation_level] HgtSIM generates as output the mutated gene inserted into the recipient genome, along with its corresponding translated protein sequence. Given that the tree feature-extraction pipeline relies on NCBI protein accession identifiers to retrieve taxonomic metadata, and the mutated protein sequences do not exist in the NCBI database, each simulated sequence was assigned a NCBI protein accession identifier of the corresponding recipient species to enable taxonomic annotation during the subsequent phylogenetic analysis.

To obtain a comprehensive set of homologous sequences for downstream phylogenetic analysis, protein similarity searches were performed using DIAMOND v2.1.9, using the transferred protein as the query against the NCBI NR database. For simulations representing the HGT, Multiple MC, and Patchy phylogeny patterns, searches were restricted to prokaryotic sequences using the parameter --taxonlist 2,2157, corresponding to Bacteria (taxid 2) and Archaea (taxid 2157). In contrast, for the Limited donor evidence and NoHGT patterns, searches were restricted to fungal sequences using the parameter --taxonlist 4751, corresponding to Fungi taxonomic identifier.

Homology searches results were filtered to retain only high-confidence homologs. Hits were excluded if they exhibited a pairwise identity below 30%, query coverage below 60%, or an E-value greater than 1 × 10⁻⁵. The amino acid sequences were retrieved using *efetch* and combined with the mutated protein sequences into a single FASTA file.

For each simulation, the complete set of homologous sequences was processed through a standardized phylogenetic reconstruction workflow. Multiple sequence alignments were generated using MAFFT v7.520 (75) under the automatic parameter selection mode. Ambiguously aligned regions were filtered out using Gblocks v0.91b (76) to reduce alignment noise. The curated alignments were then converted to PHYLIP format, and the most appropriate amino acid substitution model for each dataset was identified using ModelTest-NG v0.1.7 (77).

Maximum-likelihood phylogenies were inferred with PhyML v3.3.20220408 (78), and branch support was evaluated using Shimodaira–Hasegawa (SH)–like approximate likelihood ratio test values. The resulting trees were exported in NEXUS format and annotated using a custom Python script v3.10.12, which integrated taxonomic metadata retrieved from the corresponding GenBank records for all sequences included in each phylogeny.

### Generation of phylogenetic trees from published iHGT candidates

To evaluate model performance under real biological conditions, we compiled iHGT candidate genes from Aguirre-Carvajal & Armijos-Jaramillo (2026) and Aguirre-Carvajal et al. (2025). Prior to analysis, it was verified that none of these candidates had been previously used during model training. Redundant entries between the two studies were removed, resulting in a final dataset of 1,438 HGT candidates. Phylogenetic trees for the candidates reported by Aguirre-Carvajal & Armijos-Jaramillo (2026) were retrieved from the corresponding public repository (https://doi.org/10.5281/zenodo.18435322).

For candidates studied by Aguirre-Carvajal et al. (2025), phylogenetic trees were reconstructed. First, protein similarity searches were performed using DIAMOND, with the candidate protein used as the query against the same reference database described above. To ensure comprehensive taxonomic representation, two separate searches were conducted: one restricted to prokaryotic sequences (--taxonlist 2,2157) and another restricted to eukaryotic sequences (--taxonlist 2759, corresponding to Eukaryota). The resulting hits were merged and filtered following the criteria described previously and subsequently processed through the phylogenetic reconstruction pipeline described in the previous section.

The inferred phylogenetic trees were manually inspected using Geneious Prime 2025.2.1 (https://www.geneious.com; accessed February 26, 2026) and classified according to the phylogenetic patterns defined in Table 3. Phylogenetic trees from Aguirre-Carvajal & Armijos-Jaramillo (2026) were assigned the same phylogenetic patterns as reported in the original study.

### Application of AVP for phylogenetic analysis

AVP (Alienness vs. Predictor) v1.0.10 (28) was used as a benchmark method to evaluate the performance of the proposed approach. AVP was selected for comparison because it represents one of the few automated methods specifically designed to detect iHGT events using phylogenetic tree topology. Like the approach developed in this study, AVP relies on phylogenetic tree topology to infer evolutionary scenarios, making it conceptually comparable for evaluating classification performance. Candidates were analyzed using AVP in IQ-TREE mode to systematically assess the inferred phylogenetic topologies for each case. Based on these patterns, AVP classified each candidate as HGT or NO. AVP may also assign inconclusive classifications (e.g., Unknown or Complex).

AVP requires as input the candidate query sequence, the corresponding homology search results, and two taxonomic parameters: the ingroup and the exclusion group parameter (EGP). For simulated datasets, the mutated homologues generated by HgtSIM are absent from public databases. To include them in the homology search results required by AVP, a custom DIAMOND reference database was constructed for each simulation using the mutated sequences. The transferred protein was subsequently used as the query to compute homology scores against this custom database. These scores were then merged with the general homology search results prior to AVP analysis.

In this framework, the ingroup represents the focal taxonomic lineage used to detect potential external donors, whereas the EGP defines the taxonomic group containing the candidate sequence and specifies the lineage in which horizontal transfer is hypothesized to have occurred. The ingroup and EGP definitions applied in each analysis are provided in Supplementary Table 4. Detailed instructions for installing and running AVP are available in the AVP GitHub repository (https://github.com/GDKO/AvP).

### Comparative benchmarking of iHGT detection methods

Phylogenetic trees derived from both simulated datasets (1000 trees) and real biological candidates (1438 trees) were processed through the feature extraction pipeline, and the resulting feature matrices were used as input for the trained classification model to assign each case to one of the predefined phylogenetic patterns. The parameters used to run the pipeline are summarized in Supplementary Table 5.

The classifications produced by AVP and by the developed model were systematically compared. For both the simulated dataset and the real biological dataset, the curated labels were used as the reference classification. To ensure comparability between the two tools, the phylogenetic conclusions were mapped to the categories defined in Table 3. In the case of AVP, internal classifications that did not correspond to either iHGT or NoHGT were grouped into the Inconclusive category.

The predictions generated by the Random Forest (RF) model and by AVP were evaluated against the reference classification using several performance metrics. Accuracy was calculated as the proportion of correctly classified instances. In addition, precision, recall, and F1-score were computed using macro-averaging across the three classes (*iHGT*, *NoHGT*, and *Inconclusive*) to ensure that each class contributed equally to the final metric regardless of class frequency. These metrics were calculated using the confusionMatrix function implemented in the caret package v6.0-93 in R.

To further assess classification quality, the Matthews Correlation Coefficient (MCC) was also calculated using the mcc function from the mltools package v0.3.5. MCC provides a balanced measure of classification performance by incorporating true and false positives and negatives, making it particularly suitable for evaluating multi-class classification problems. All metrics were computed in the R statistical environment using confusion matrices derived from the predicted and reference labels.

All software tools used in this study are freely available and open-source, unless otherwise specified. Python and R based analyses were conducted using publicly available libraries, and all external tools used for sequence analysis and phylogenetic reconstruction (e.g., DIAMOND, MAFFT, PhyML, and HgtSIM) are freely accessible for academic use. Manual inspection of phylogenetic trees was performed using Geneious Prime.

To facilitate interpretation of the complete analytical framework, a global overview of the methodological workflow is provided in Figure 9. This diagram summarizes the sequence of computational steps, software tools, and data transformations involved from data acquisition to model evaluation and benchmarking.

**Figure 9.**
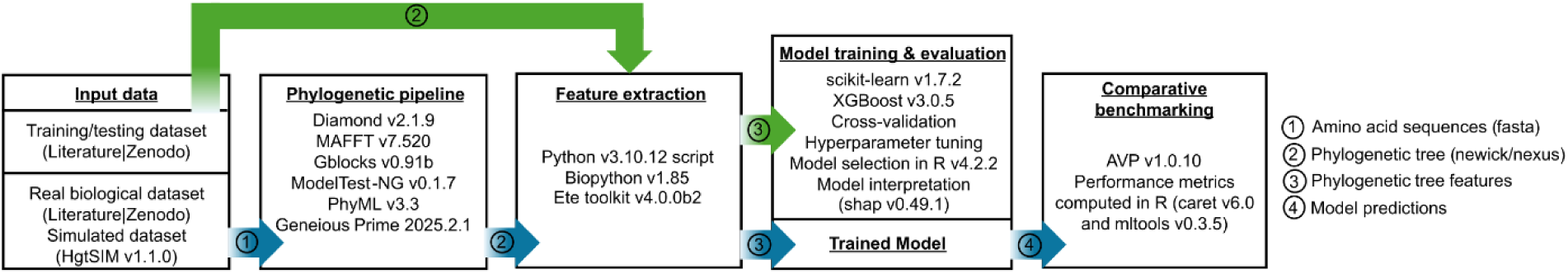
Computational workflow of the proposed framework for phylogenetic pattern classification and benchmarking. The workflow begins with three sources of input data: (i) a curated training/testing dataset, (ii) real biological datasets, and (iii) simulated datasets generated with HgtSIM. Protein sequences are processed through a phylogenetic reconstruction pipeline to generate gene trees. These trees are subsequently used for feature extraction, where taxonomic metadata and phylogenetic descriptors are computed using Python-based tools. The resulting feature matrices are used to train and evaluate machine-learning models, including cross-validation, hyperparameter tuning, model selection and interpretation. The best-performing model is then applied to classify phylogenetic patterns, and its predictions are benchmarked against AVP. Performance metrics are computed in R. Green arrows represent the training and model development workflow, whereas blue arrows indicate the benchmarking and evaluation workflow applied to independent datasets. Numbered arrows indicate data types transferred between steps: (1) amino acid sequences, (2) phylogenetic trees, (3) extracted features, and (4) model predictions.

Each stage of the workflow is described in detail in the preceding sections of the methodology. Taken together, this integrated framework provides a coherent and reproducible overview of the procedures implemented to train and evaluate the classification model for phylogenetic pattern identification, as well as to systematically compare its performance with that of alternative approaches such as AVP.

The complete implementation of the proposed pipeline, including feature extraction, model training, prediction, and benchmarking procedures, is publicly available in the project’s GitHub repository (https://github.com/kevynaguirre/PhyloHGT). All training and evaluation workflows are fully documented to ensure transparency and reproducibility, including detailed scripts. In addition, phylogenetic trees from both the simulated datasets and the unseen real biological candidates used in this study have been deposited in the same repository, enabling full replication of the analyses.

## Conclusion

In this study, we present a reproducible and interpretable framework for classifying phylogenetic patterns associated with interkingdom horizontal gene transfer (iHGT). By formalizing expert-driven phylogenetic interpretation into a supervised machine-learning problem, we demonstrate that complex evolutionary scenarios can be systematically represented using a small set of biologically meaningful features derived from gene tree topology and taxonomic composition.

Our results show that topological relationships—particularly the relative position of additional lineages within the tree—constitute the most informative signal for distinguishing among evolutionary scenarios. By integrating these signals into a multi-class classification framework, the proposed approach moves beyond traditional binary detection strategies and explicitly accounts for ambiguous cases, providing a more realistic representation of evolutionary complexity.

The framework achieved high predictive performance across both simulated and real biological datasets and consistently outperformed existing approaches, highlighting the advantages of incorporating global phylogenetic structure into automated inference. At the same time, the analysis underscores that certain phylogenetic patterns commonly interpreted as evidence of iHGT may arise from alternative evolutionary processes, emphasizing the need for cautious and context-aware interpretation.

Beyond improving iHGT detection, this work establishes a general strategy for translating qualitative phylogenetic reasoning into quantitative, scalable, and reproducible models. As genomic datasets continue to expand, such approaches will be essential for enabling robust evolutionary inference and for developing standardized criteria in comparative genomics.

## Authors contribution

Conceptualization: K.A.-C., V.A.-J.; Methodology: K.A.-C., V.A.-J.; Software: K.A.-C.; Validation: K.A.-C., C.R.M.; Formal analysis: K.A.-C., V.A.-J.; Investigation: K.A.-C., V.A.-J.; Data curation: K.A.-C., V.A.-J.; Resources: V.A.-J., C.R.M.; Supervision: V.A.-J., C.R.M.; Project administration: V.A.-J., C.R.M.; Funding acquisition: V.A.-J., C.R.M.; Writing – original draft: K.A.-C., V.A.-J.; Writing – review & editing: K.A.-C., V.A.-J, C.R.M.

## Data availability statement

All data and code used in this study are publicly available. The complete implementation of the PhyloHGT pipeline, including scripts for feature extraction, model training, prediction, and benchmarking, is available in the GitHub repository: https://github.com/kevynaguirre/PhyloHGT

The repository also includes the datasets generated and analyzed during this study, including feature matrices, simulated phylogenetic trees, and phylogenetic trees derived from real biological candidates. All files necessary to reproduce the analyses and results are provided within the repository and Supplementary Tables.

